# Bacterial extracellular vesicles exhibit distinct functional potential across the biogeographic provinces of the South Pacific Ocean

**DOI:** 10.1101/2025.06.30.662282

**Authors:** Eduard Fadeev, Neza Orel, Tinkara Tinta, Leila Afjehi-Sadat, Haoran Liu, Thomas J. Browning, Zhongwei Yuan, Eric P. Achterberg, Steven J. Biller, Daniel J. Sher, Gerhard J. Herndl

## Abstract

Bacterial extracellular vesicles (BEVs) are nanoscale membranous structures released by diverse types of bacteria. Laboratory model systems indicate that these nanoparticles may play several roles in the ecophysiology of marine bacteria. However, their actual functionality in the environment remains unclear. Here we describe the proteomic composition of marine BEVs over more than 5,000 nautical miles of surface waters in the South Pacific, linking BEV cargoes to the bacterial communities producing them. BEVs were consistently present across a range of biogeochemical conditions, with an overall abundance comparable to that of bacterial cells. However, the protein content of the BEVs varied significantly between different ocean regions. The BEVs were enriched in carbohydrate transporters under phytoplankton bloom conditions, and contained iron and phosphate uptake-related proteins in nutrient-limited waters. This suggests that BEVs could enable cells to perform key extracellular functions in the marine environment. Our observations further highlight the prevalence of BEVs and the biogeographic patterns of their functional potential across oceanic scales.

## Introduction

Bacterial extracellular vesicles (BEVs) are nanoscale, membrane-bound structures typically ranging in diameter from 20 to 250 nm, which are released by bacterial cells. They can result from ‘blebbing’ mechanisms in intact cells or be formed as a by-product of cell lysis. This can influence their contents and, consequently, their functional potential (Kulp & Kuehn, 2010). Since their discovery in human-associated *Escherichia coli* in the 1960s (Knox et al., 1966), most studies of BEVs have examined their roles within laboratory model organisms, particularly host-associated bacteria in animal and plant systems. BEVs can serve as a mechanism for removing harmful compounds from the cell, but can also transport various biological cargoes (e.g., nucleic acids and proteins) over long distances in a protected and locally concentrated manner (McMillan & Kuehn, 2021). One of the best-documented functions of BEVs in intercellular interactions is the transfer of DNA molecules (Bitto et al., 2017; Soler & Forterre, 2020), which can facilitate the exchange of antibiotic resistance genes between different bacterial lineages (Yaron et al., 2000), as well as the transfer of entire plasmids (Johnston et al., 2023; Renelli et al., 2004), recently termed ‘vesiduction’ (Soler & Forterre, 2020). BEVs can also play roles in bacterial quorum sensing (Mashburn & Whiteley, 2005; Toyofuku et al., 2017), biofilm formation (Yonezawa et al., 2009), and contribute to microbial “warfare” by delivering compounds that kill competing bacteria (Kadurugamuwa & Beveridge, 1996; Z. Li et al., 1998). As BEVs can contain a variety of components released by bacterial cells, including enzymes, receptors and transporters derived from cell membranes, they may interact with organic and inorganic chemicals in the environment (Orench-Rivera & Kuehn, 2016; Zlatkov et al., 2020). Such extracellular activities, including the enzymatic digestion or binding of various environmental compounds, are particularly important for bacteria living in aqueous environments (Toyofuku et al., 2023, 2015).

The ocean, where bacteria are among the most abundant organisms (Bar-On & Milo, 2019), is a highly dynamic environment both physically and biochemically. The surface ocean is characterised by a depletion of bioavailable nutrients required for life in some vast areas (e.g., oligotrophic oceanic gyres) (Browning et al., 2017) and a high availability in others (e.g., coastal upwelling regions) (Capone & Hutchins, 2013; Moore et al., 2013). To survive in such ecosystems, marine bacteria have developed a wide range of ecophysiological capabilities (Stocker, 2012; Worden et al., 2015; Zoccarato et al., 2022), and implement highly diverse nutrient and energy utilisation strategies (Kirchman, 1994; Reintjes et al., 2017; Tortell et al., 1999). In the interaction between marine bacteria and their oligotrophic environment, high cellular surface area to volume ratios facilitate efficient nutrient uptake (Gasol et al., 1995; Koch, 1996). In this sense, the release of marine BEVs by bacterial cells can be conceptualised as an expansion of the cell surface which disperses through the aqueous medium (Kulp & Kuehn, 2010), randomly encountering various components of the marine environment. Laboratory observations of several isolates of globally abundant marine bacteria suggest that BEVs may mediate specific extracellular functions in the ecophysiology of these bacteria, such as binding nutrients (e.g., iron or phosphate) and delivering them in concentrated form to bacterial cells (Biller et al., 2023; Biller, Lundeen, et al., 2022; Biller et al., 2014; Dürwald et al., 2021; Fadeev et al., 2023; Fischer et al., 2019). However, despite the potential importance of BEVs for marine bacteria, and ubiquity in the marine environment (Linney et al., 2022; Lücking et al., 2023), our understanding of their biogeographic diversity across large environmental gradients is unknown.

To address this knowledge gap, we conducted a spatially extensive sampling to examine the abundance and protein cargo diversity of BEVs across the South Pacific Ocean. We determined how the molecular cargo of marine BEV populations varies in context with their local microbial and biogeochemical environment, finding that BEVs may play distinct functions for different microbial populations. Our observations across large oceanic regions reinforce the emerging picture that important extracellular bacterial functions may be mediated by BEVs and can inform future efforts to test specific hypotheses about their ecological roles.

## Results and Discussion

### Distinct biogeochemical conditions across the South Pacific Ocean

To investigate the potential ecophysiological functions of BEVs in marine bacterial communities we sampled across diverse environmental conditions, varying in nutrient limitations and phytoplankton growth conditions. We collected surface water (5-6 m depth) at 20 stations along a basin-scale transect that spanned the South Pacific Ocean (Fig. 1; Table S1). A detailed physicochemical characterisation of this transect showed that field sampling was carried out across four oceanic provinces (Liu et al., 2024; Yuan et al., 2025). Briefly, the easternmost region was the Chilean coastal upwelling zone (UP), characterized by elevated nitrate and phosphate concentrations, but depleted in dissolved iron. In this area there were elevated chlorophyll-a concentrations (ca. 3 mg m^-3^; Fig. S1), with the phytoplankton community mostly composed of diatoms (Fig. 1). The transition zone between the upwelling and the subtropical gyre (TRAN) was characterized by elevated concentrations of phosphate and dissolved iron, but was depleted in nitrate (Fig. S1). This area also had relatively elevated chlorophyll-a concentrations (0.04-0.26 mg m^-3^), with mixed phytoplankton communities. The centre of the subtropical gyre (GYRE) was characterized by very low nitrate concentrations and a declining spatial gradient from east to west of phosphate concentrations, with strongly depleted chlorophyll-a concentrations (< 0.03 mg m^-3^; Fig. S1). The westernmost region (WEST) was defined as the area of the subtropical gyre with the lowest phosphate concentrations and higher (compared to GYRE) chlorophyll-a concentrations (0.03-0.14 mg m^-3^; Fig. S1), with the phytoplankton comprising predominantly cyanobacteria.

**Fig. 1.**
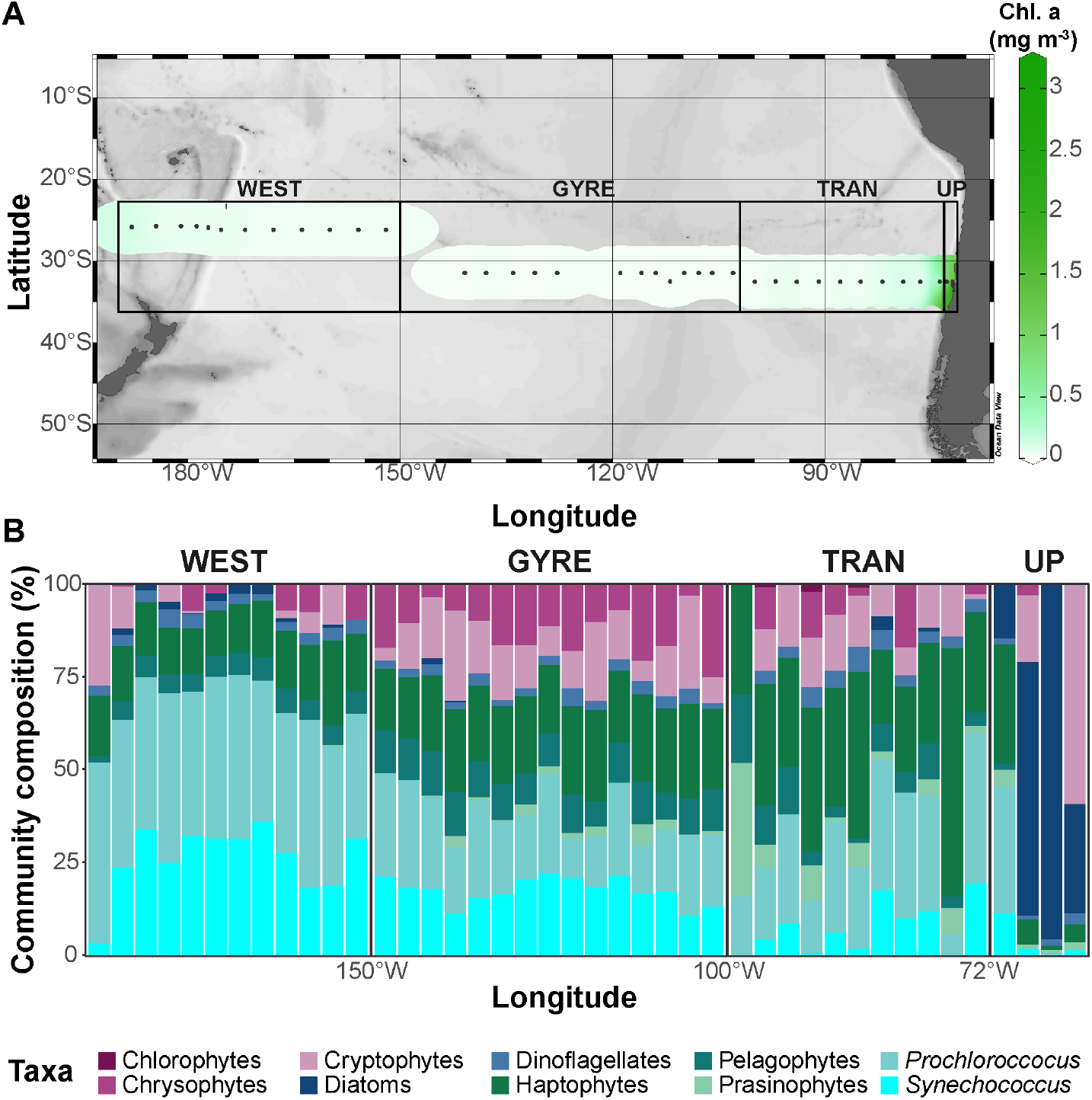
Diverse biogeochemical regimes encountered across the South Pacific Ocean. (A) Spatial grouping of the sampling stations according to four oceanic provinces: ‘UP’ - Chilean coast upwelling zone; ‘TRAN’ - transition zone between the upwelling and the subtropical gyre; ‘GYRE’ - South Pacific subtropical gyre; ‘WEST’ - westernmost region. (B) The composition of the surface water phytoplankton at each station, estimated on the basis of diagnostic pigments.

We found that bacterial abundance varied across the biogeochemical provinces (Table S1), and the taxonomic composition of the bacterial communities (Fig. 2) also differed significantly between them (PERMANOVA test; F_3,14_=21.16, R^2^=0.82, p=0.001). To investigate the physiological and metabolic processes expressed within the bacterial communities, we characterized the metaproteome of the bacterial cells, identifying between 4281-6200 cellular proteins at each station (Table S2). The composition of bacterial cellular proteins significantly varied between the four regions (PERMANOVA test; F_3,16_=3.06, R^2^=0.36, p=0.001; Fig. S2) and further differed between each pair of adjacent provinces (Table S3), indicating that these represented distinct functional ecosystems. Bacterial cell abundance was the highest in the upwelling zone (9-26 ×10^8^ cells L^-1^), with the bacterial community composition showing a strong heterotrophic response to a diatom bloom (Fig. 2). This response was attributed mainly to members of the classes Gammaproteobacteria and Flavobacteria, which are typically active during elevated availability of organic carbon (Pontiller et al., 2022; Teeling et al., 2016). In the transition zone, bacterial abundance reached 0.7-3.5 ×10^8^ cells L^-1^, with the cyanobacteria *Prochlorococcus* and *Synechococcus* comprising more than 50% of the bacterial community (Table S1). The elevated photosynthetic cyanobacterial activity in this zone was potentially stimulated by a nutrient influx from a returning lateral current (Liu et al., 2024). The subtropical gyre had the lowest bacterial cell abundances (0.2-0.9 ×10^8^ cells L^-1^), with cyanobacteria comprising up to 50% of the bacterial community (Table S1). Metagenomic characterization of the bacterial community composition further suggested dominance of the oligotrophic Alphaproteobacterial order *Pelagibacterales* (Fig. 2)(R. M. Morris et al., 2002). This lineage has previously been shown to numerically dominate the bacterial communities of the ultra-oligotrophic South Pacific subtropical gyre (Oggerin et al., 2024; Reintjes et al., 2019). In the westernmost region, cell abundance was significantly higher than in the subtropical gyre (0.8-1.3 ×10^8^ cells L^-1^; Student’s t-test, p=0.002), with a predominance of *Prochlorococcus*. The photosynthetic cyanobacterial activity in this region may have been partly facilitated by input from a large volcanic eruption in the region two months prior to sampling (Zhang et al., 2024).

**Fig. 2.**
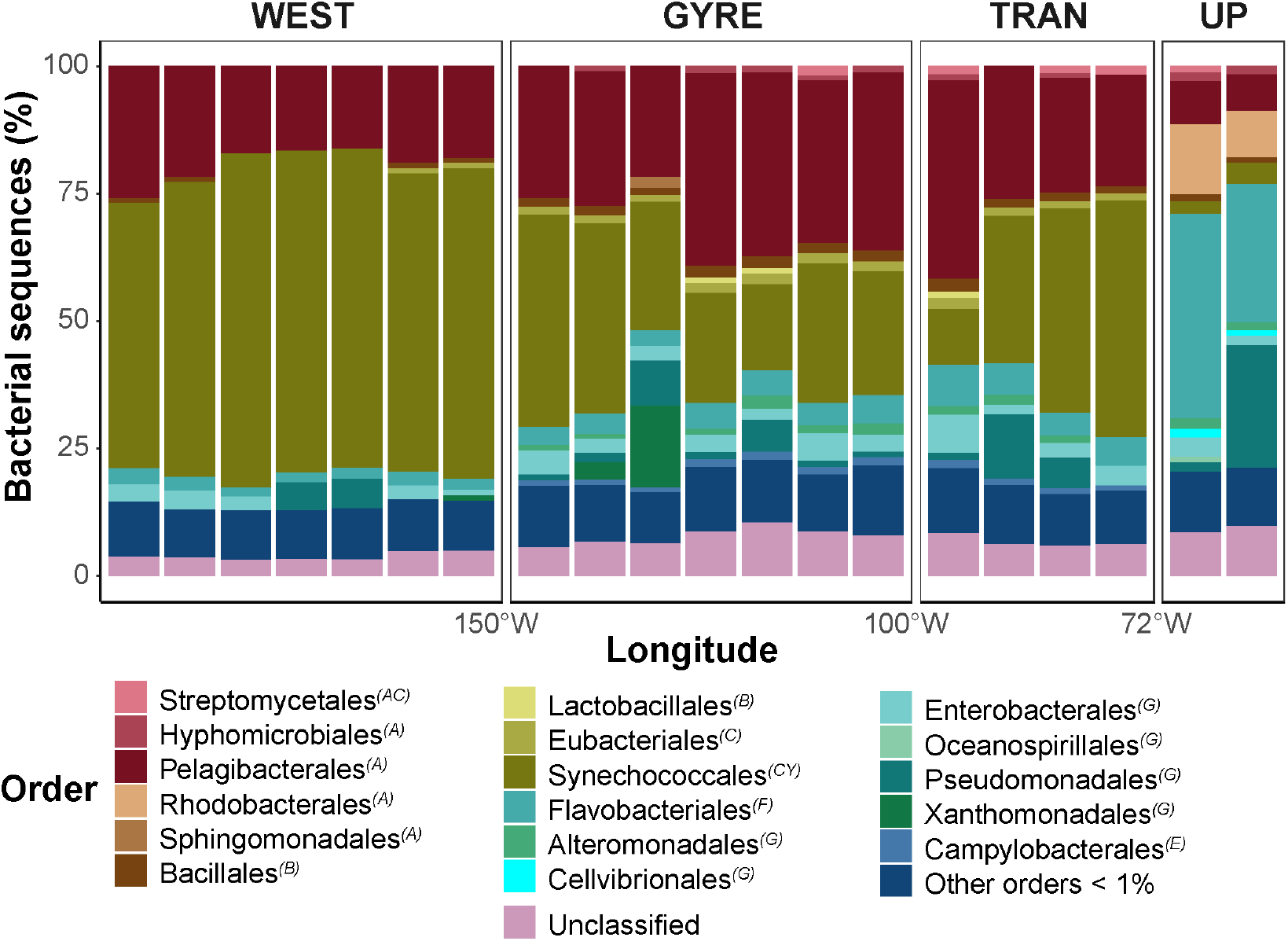
Bacterial community composition based on taxonomic classification of metagenomic sequences. The taxonomic class of each order is given in parentheses: ‘AC’ – Actinomycetes, ‘A’ – Alphaproteobacteria, ‘B’ – Bacilli, ‘C’ – Clostridia, ‘CY’ – Cyanophyceae, ‘F’ – Flavobacteriia, ‘G’ – Gammaproteobacteria, ‘E’ – Epsilonproteobacteria. ‘Other orders < 1%’ – refers to taxonomic orders that comprised less than 1% of the sequences; ‘unclassified’ – refers to bacterial sequences with unknown classification on an order level. All samples contained 2-5×10^8^ classified sequences.

### Abundance, characteristics, and main producers of marine BEVs

To investigate the abundance and diversity of marine BEVs, we sampled the extracellular size fraction corresponding to BEV-like structures (100 kDa-0.22 µm) across all stations using size fractionated filtration. The size distribution of marine viruses overlaps with that of BEVs (Biller et al., 2017; Nekrouf et al., 2025; Soler et al., 2015), so we used density gradient separation to further enrich our samples for BEVs in the laboratory (see Materials and Methods). At least 50% of the purified BEVs were between 70-130 nm in diameter (Fig. 3), matching the size distributions previously observed in cultures of marine bacteria (Biller et al., 2023, 2014). Their abundance was overall comparable to the abundance of bacterial cells (Fig. 3), although this is likely a conservative estimate, since the density gradient separation led to a reduced yield of 6-52% of all nanoparticles (Fig. S3), and the storage of the samples on board at -20°C may also have resulted in a loss of BEV-like structures (Ahmadian et al., 2024).

**Fig. 3.**
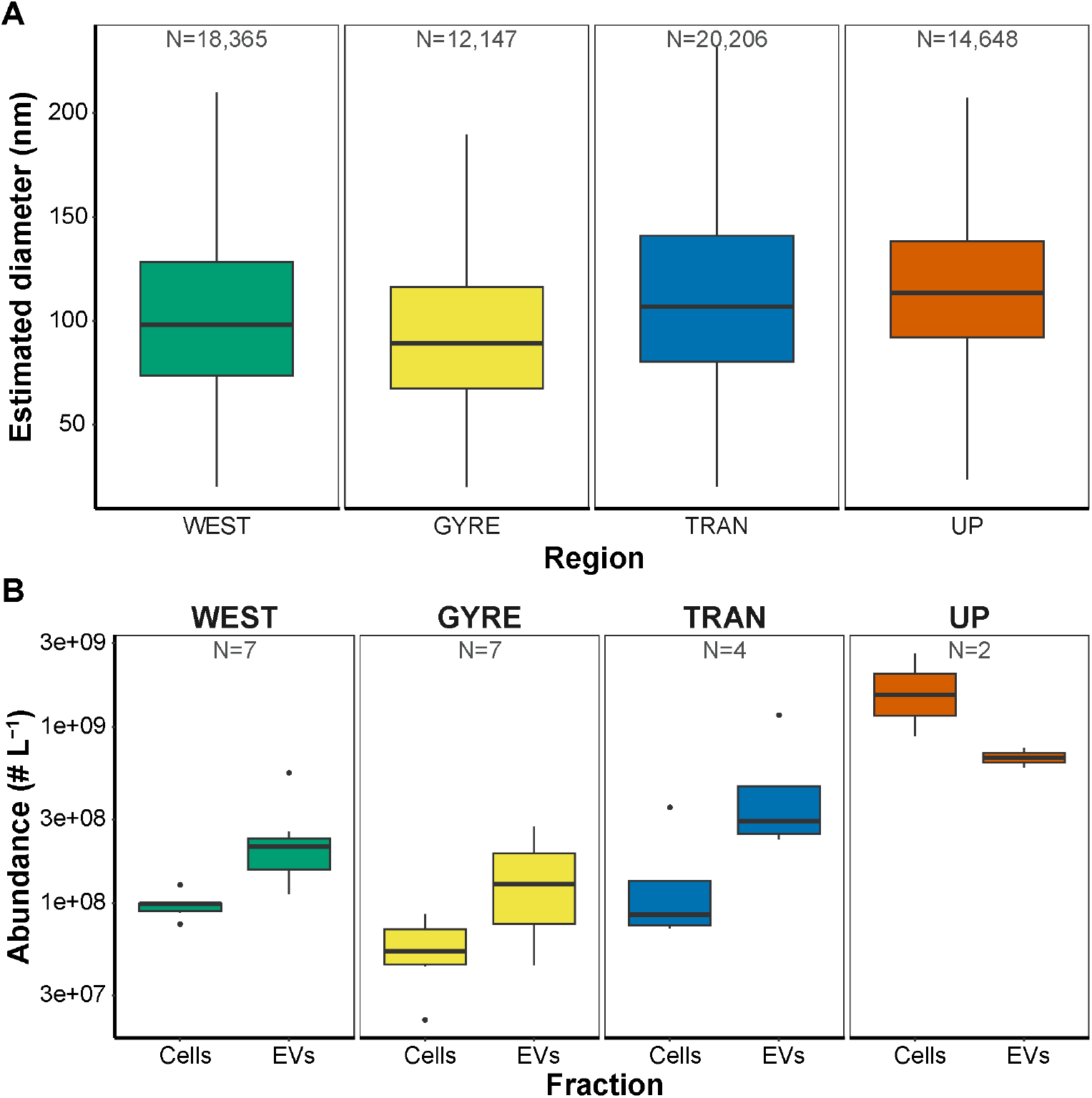
Physical characteristics of marine BEV-like structures. (A) Estimated diameter distribution of the BEV-like structures. The error bars represent variation between NTA technical replicates. (B) Bacterial cell and BEVs abundances, estimated using flow cytometry and nanoparticle tracking analysis (NTA), respectively.

We then characterised the bulk protein content in the BEV-designated density fraction (see Materials and Methods) and identified 1940-3394 proteins at each station (Table S1). Of these proteins only ca. 3% (58-123 proteins) were taxonomically or functionally (e.g., capsid or tail sheath) related to viruses, suggesting a strong enrichment of BEVs over other types of protein-containing marine nanoparticles in our samples. We found that 78-96% of proteins observed in the BEVs were also found in the corresponding cellular fraction at each station, but the composition of the proteins significantly varied between the two fractions (PERMANOVA test; F_1,38_=15.01, R^2^=0.28, p=0.001). This indicated that the BEV-associated proteins represented a distinct subset of the bacterial cellular proteome.

Marine bacteria, which are mostly Gram-negative, can produce BEVs through blebbing of the outer membrane, leading to preferential incorporation of outer membrane and periplasmic material from the releasing cell (Kulp & Kuehn, 2010; Schwechheimer et al., 2013). External conditions have been shown to have an impact on the release of BEVs, potentially leading to some degree of preferential protein packaging (Gerritzen et al., 2018; Minhas & Greenman, 1989; Orench-Rivera & Kuehn, 2021; Sabra et al., 2003). Other BEVs may result from bacterial cell lysis, including due to bacteriophages, and thus represent a random assemblage of proteins from different subcellular origins (Pérez-Cruz et al., 2015; Toyofuku et al., 2023). To differentiate between these different origins of BEVs, we determined whether there is an enrichment in non-cytoplasmic proteins in the BEV fraction. We carried out a protein enrichment analysis on all samples in the BEV fraction as compared with the cellular fraction, and identified a total of 1065 proteins that were significantly enriched in the BEVs (absolute log_2_ fold difference > 1, adjusted p-value < 0.1), relative to their abundance in the cells. By comparison, 2017 proteins were significantly enriched in the cellular fraction (Table S4). Prediction of sub-cellular origins revealed that half of the BEV-enriched proteins were non-cytoplasmic, comprising mostly outer membrane and extracellular proteins, compared to ca. 25% of such proteins enriched in the cellular fraction (Fig. 4). Although we cannot exclude the presence of lysis-derived BEVs in our samples, the enrichment of mostly non-cytoplasmic proteins suggests that the marine BEVs in our samples may have derived primarily from membrane blebbing of bacterial cells (Toyofuku et al., 2023).

**Fig. 4.**
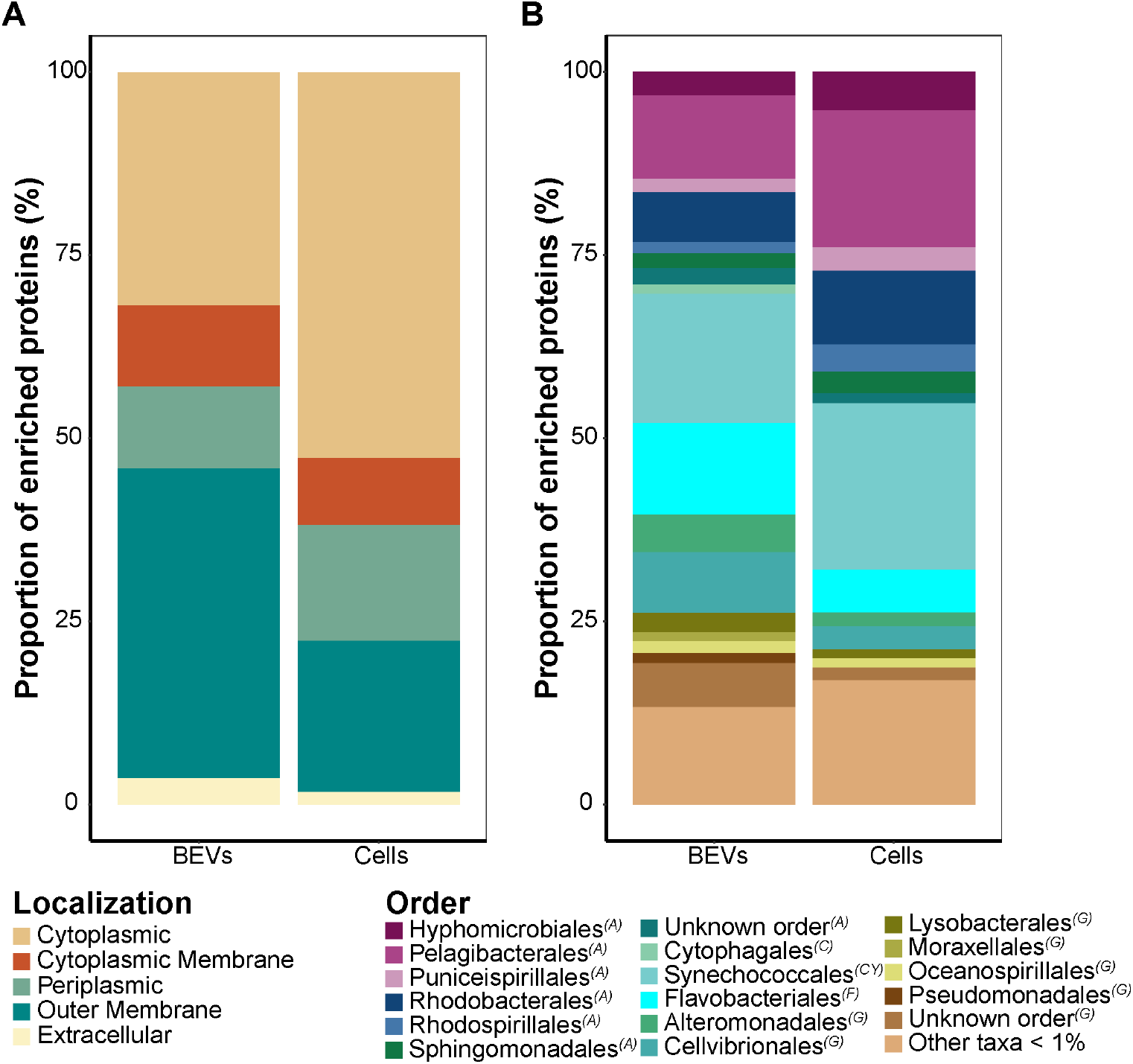
Overview of significantly enriched proteins in BEVs and cellular fractions. The bar plots represent the composition of all significantly enriched proteins according to their estimated sub-cellular localization (A) and taxonomic origin (B). The total number of enriched proteins in the BEVs and the cellular fractions was 1121 and 2075, respectively. Taxonomic composition of enriched proteins in the cellular and BEVs fractions according to different oceanic provinces. The taxonomic class of each order is given in parentheses: ‘A’ – Alphaproteobacteria, ‘C’ – Cytophagia, ‘CY’ – Cyanophyceae, ‘F’ – Flavobacteriia, ‘G’ – Gammaproteobacteria.

Studies of laboratory cultures indicate that the major classes of marine bacteria exhibit significant variations in BEV production rates (Biller et al., 2023). Our protein enrichment analyses from field samples revealed that the two taxonomic lineages with the largest number of enriched proteins in both the BEVs and in the cellular fractions were the orders *Synechococcales* (including *Prochlorococcus* and *Synechococcus*) and the order *Pelagibacterales* (class Alphaproteobacteria). The ratio of enriched proteins between the BEVs and cellular fractions of these lineages showed a clear tendency towards the latter (Fig. 5). In contrast, enriched proteins associated with other bacterial orders, such as the *Alteromonadales* and *Cellvibrionales* (class Gammaproteobacteria), and the order *Flavobacteriales* (class Flavobacteriia), accounted for more than twice BEV-enriched proteins compared to the cellular fraction (Fig. 5). These observations could indicate that BEVs from these lineages contain more protein on average than those from Alphaproteobacteria and cyanobacteria. Alternatively, based on the assumption that the number of enriched proteins reflects to some extent the abundance of BEVs, these results are consistent with the previously raised hypothesis that fast-growing heterotrophic marine bacteria contribute more to the BEV population than slow growing ones (Biller et al., 2023).

**Fig. 5.**
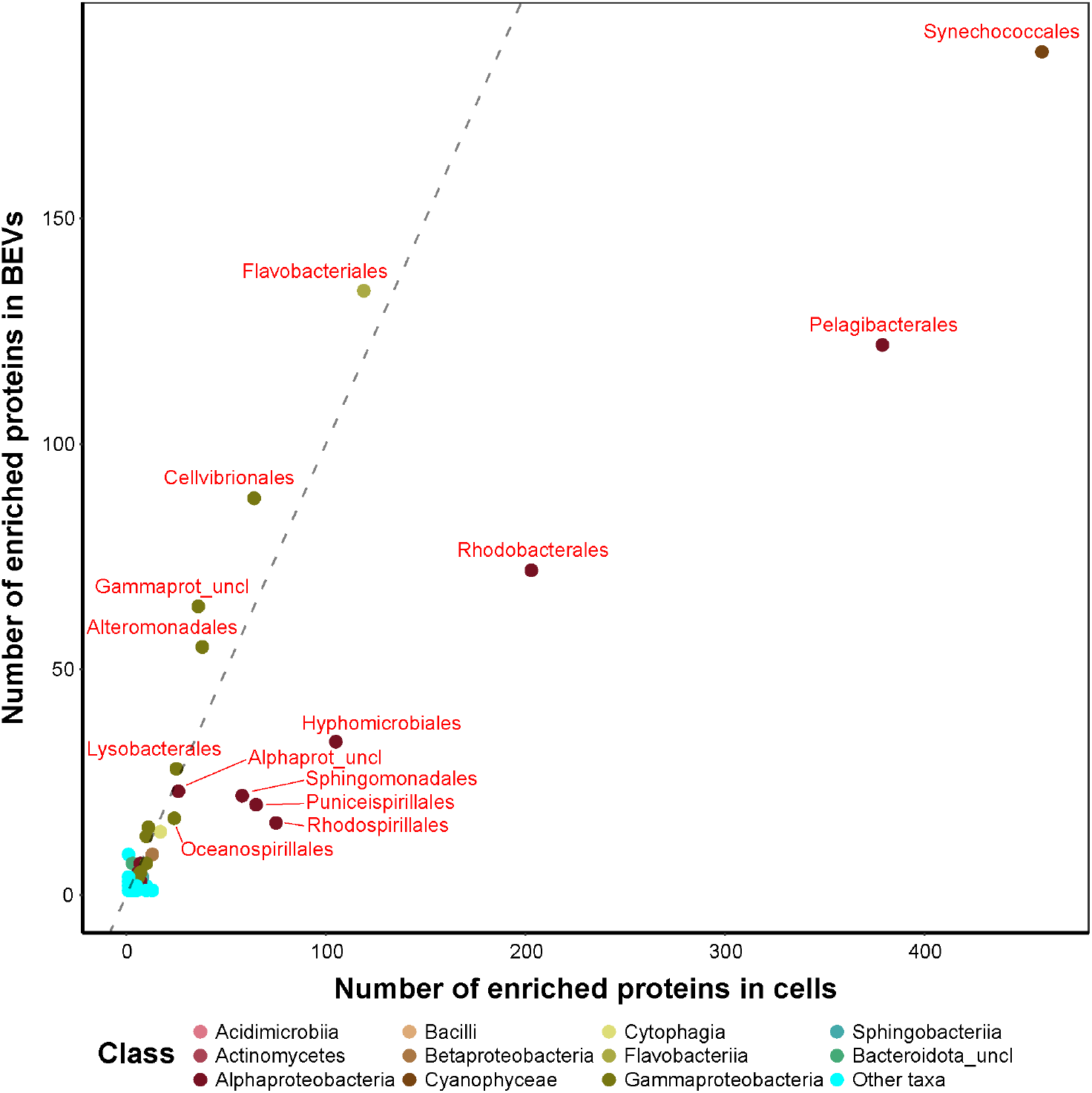
Ratio of significantly enriched proteins in BEVs and cellular fraction in each taxonomic order. The dashed line represents a theoretical ratio of 1:1 between the number of proteins enriched in each fraction. ‘Other taxa’ represents classes with less than 3 enriched proteins in each of the fractions.

The distinct biogeochemical conditions sampled in this study shape bacterial communities (Fig. 2), which could potentially affect BEV populations and their functions (Biller et al., 2023; Orench-Rivera & Kuehn, 2016). Therefore, in order to infer the potential functions of BEVs while accounting for these differences, we examined the proteins enriched in BEVs separately for each oceanic province. As only two stations sampled in the upwelling region, this region was excluded from further analysis. We found a total of 1295 proteins (Table S5), which were associated with 77 different bacterial orders (Fig. S4), that were significantly enriched in BEVs compared to bacterial cells. Different functional groups of proteins were found enriched in BEVs in different oceanic provinces, suggesting that they may play a role in multiple ecophysiological processes, the main ones of which are discussed below.

### Marine BEVs as mediators for extracellular electron transfer?

The surface waters of the South Pacific receive high solar irradiance. High light levels with associated enhanced UV radiation can induce oxidative stress in photosynthetic microorganisms (Mella-Flores et al., 2012). Marine cyanobacteria (e.g., *Prochlorococcaceae* and *Synechococcaceae*) have developed

various strategies to cope with this environmental stressor, one of which is the transfer of excess electrons to ferredoxins and flavoproteins (Scanlan et al., 2009). In our dataset, we observed that 10-30% of all enriched cyanobacterial proteins in the BEV fraction were outer-membrane porins, compared to 0.5-5% in the cellular fraction (Table S5). We also found ferredoxins and flavoproteins, which are typically soluble, cytoplasmic proteins associated with the thylakoid membranes (Pierella Karlusich & Carrillo, 2017), in cyanobacterial BEVs across all regions, with the strongest enrichment in the centre of the subtropical gyre (Fig. 6). This enrichment under the nutrient limiting conditions may indicate that such reduced electron carriers are exported in cyanobacterial BEVs as a cellular defence mechanism against oxidative stress (Orench-Rivera & Kuehn, 2021).

**Fig. 6.**
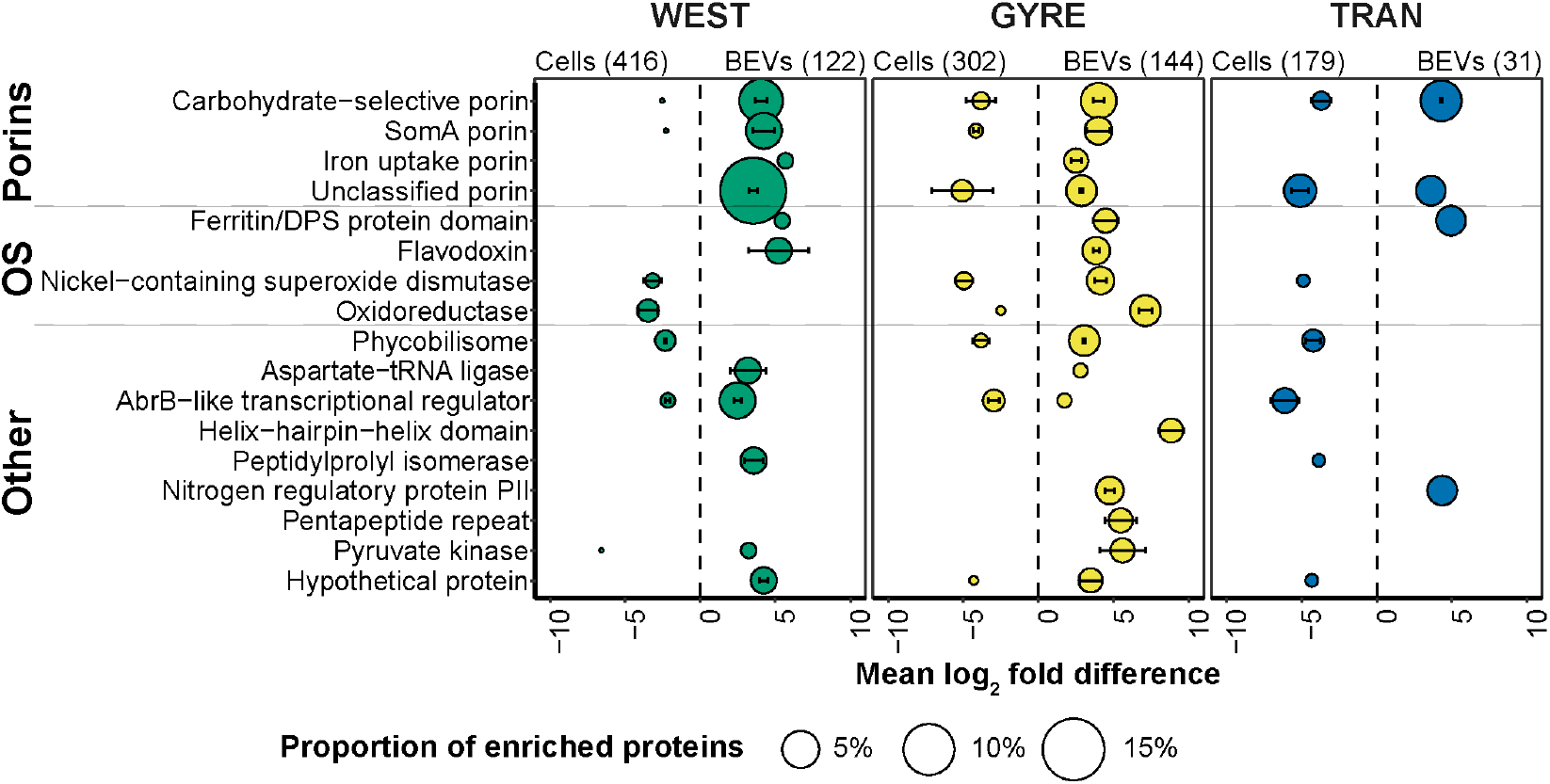
Cyanobacterial proteins significantly enriched in BEVs and cellular fractions in different regions. The mean log_2_ fold difference was calculated for each functional protein group in each fraction. Positive log_2_ fold difference represents significant enrichment in the BEVs fraction, negative mean log_2_ fold difference represents significant enrichment in the cellular fraction. The total number of enriched proteins in the BEVs and the cellular fractions is provided in brackets. The proteins were grouped according to their proposed functions in BEVs (porins, oxidative stress proteins (OS) and other proteins). For visualisation purposes, only functional groups containing at least three enriched proteins in at least one region were included.

Increasing light intensity, which can lead to oxidative stress in cyanobacteria, has been shown to increase vesiculation rates in *Prochlorococcaceae* cultures (Biller et al., 2023; Biller, Lundeen, et al., 2022). It has been also shown that marine heterotrophic bacteria, such as those belonging to the *Alteromonadales*, support the growth of *Prochlorococcaceae* by scavenging excreted compounds produced as a result of oxidative stress by the latter (J. J. Morris et al., 2011). Furthermore, *Alteromonadales* are capable of taking up BEVs produced by *Prochlorococcaceae*, suggesting that BEVs can serve as nutrient source for heterotrophic bacteria (Biller et al., 2014). Alternatively, based on the demonstrated capability of various non-marine bacteria with anaerobic capabilities to acquire energy from secreted flavoproteins (Marsili et al., 2008; Okamoto et al., 2014), we speculate that cyanobacterial BEVs enriched with electron acceptors could also serve as a direct energy source for marine heterotrophic bacteria such as the *Alteromonadales*. Given the recent observation in *Prochlorococcaceae* and *Synechococcaceae* of intercellular extensions (nanotubes), containing vesicle-like structures (Angulo-Cánovas et al., 2024), it is also possible that such extracellular electron transfer occurs through direct contact between cells. In the ocean, such extracellular electron transfer interactions are more likely to take place among particle-attached bacteria in a spatially confined environment, which is often characterized by anaerobic conditions (Bianchi et al., 2018). Although this study focuses on free-living marine bacteria, it is likely that our pre-filtration step, which aimed to remove the particulate size fraction, disrupted any such structures present in the sample. Parts of these structures may have ended up in the bulk BEVs fraction.

An alternative mechanism cyanobacteria use to cope with oxidative stress is photoprotection (Mella-Flores et al., 2012). Interestingly, while in the regions with elevated cyanobacterial activity we observed phycobilisomes enriched in the cellular fraction, in the centre of the subtropical gyre they were strongly enriched in the BEVs fraction (Fig. 6). One possible explanation is that the BEV-enriched phycobilisomes are damaged proteins that were removed from the cell (Schwechheimer & Kuehn, 2015). However, the identification of protein modifications resulting from oxidation revealed that, while cell-enriched phycobilisomes exhibited up to 5 modifications, BEV-enriched phycobilisomes exhibited only 0-1 modifications (Table S5). Therefore, it is conceivable that the release of phycobilisomes from cells into BEVs in response to light and UV stress could be a photoprotective mechanism, reducing the amount of light energy absorbed by the cell by removing light-harvesting compounds. Furthermore, given the large size of cyanobacterial phycobilisomes (6.2 MDa; Domínguez-Martín et al., 2022) and the assumption that BEVs remain attached to the cell, these phycobilisome-enriched BEVs could provide additional extracellular shading, reducing the amount of light that reaches the cyanobacterial cell.

### Marine BEVs as extracellular reserves of bio-available phosphate?

The surface waters of the South Pacific subtropical gyre are one of the most oligotrophic regions in the global ocean (Morel et al., 2010). Among the bacterial lineages that successfully inhabit such limiting environment are members of the order *Pelagibacterales* (also known as SAR11 clade). Their small cell size and large periplasm, which contains many substrate-binding proteins with exceptionally high affinities (Clifton et al., 2024; Sowell et al., 2009), which facilitates their uptake of soluble substrates (e.g., phosphate, sulphate, amino acids, and sugars) from the environment (Bosdriesz et al., 2015). This enables SAR11 to dominate the bacterial communities in the ocean (R. M. Morris et al., 2002) and thrive in nutrient-depleted marine ecosystems (Giovannoni, 2017). We observed that in all regions around one third of all proteins enriched in the cellular fraction were periplasmic substrate-binding proteins annotated to bind amino acids, compared to around 10% in the BEVs fraction, which consisted of other periplasmic binding proteins, mostly with unknown substrates (Table S5). Against this backdrop, in the centre of the subtropical gyre we found significant enrichment of five different homologues of the SAR11 phosphate binding protein PstS (Fig. 7). This periplasmic binding protein has a unique structure that has been suggested to have high affinity for phosphate, as well as being capable of binding larger organic phosphorus molecules (W.-J. Zhu et al., 2025). We propose that outer-membrane porins, which were also observed among the SAR11-derived BEV-enriched proteins (Fig. 7), form channels that allow phosphate molecules to diffuse passively into the BEVs, as they would in intact cells. They become trapped there by binding to high-affinity PstS proteins, making the BEVs a potential phosphate reservoir. Moreover, the accumulation of phosphate in the periplasm is enhanced in the presence of a proton motive force (Kamennaya et al., 2020). Therefore, the observation of proteorhodopsins enriched in BEVs of SAR11 (Fig. 7) raises the possibility that light energy could, in some individual particles, further enhance the accumulation of phosphate inside BEVs in surface waters.

**Fig. 7.**
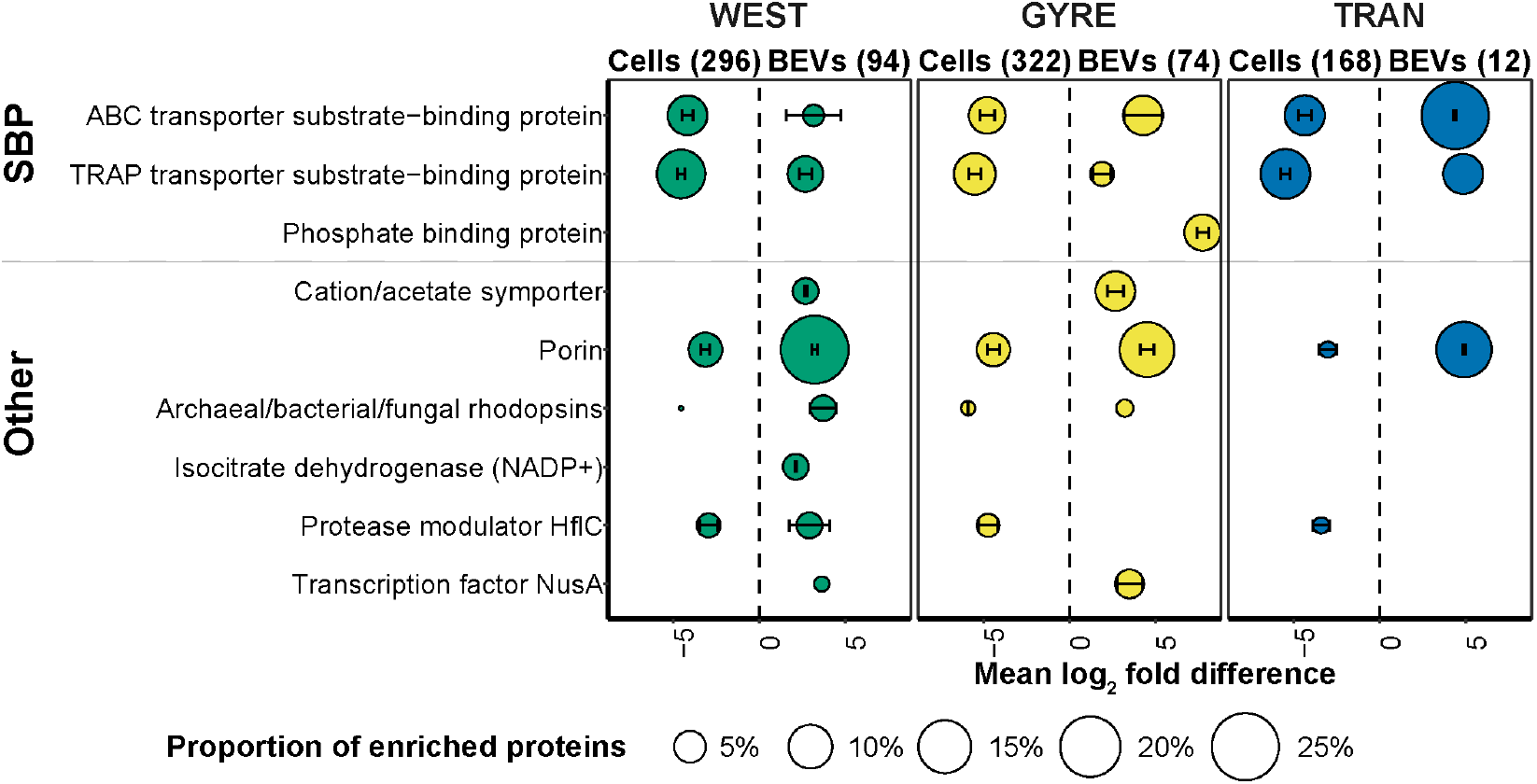
Proteins of the SAR11 clade significantly enriched in BEVs and cellular fractions in different regions. The mean log_2_ fold difference was calculated for each functional protein group in each fraction. Positive mean log_2_ fold difference represents significant enrichment in the BEVs fraction, negative mean log_2_ fold difference represents significant enrichment in the cellular fraction. The total number of enriched proteins in the BEVs and the cellular fractions is provided in brackets. The proteins were grouped according to their proposed functions in BEVs (substrate-binding proteins (SPB) and other proteins). For visualisation purposes, only functional groups containing at least two enriched proteins in at least one region were included.

### Marine BEVs as passive scavengers of iron?

Unlike the highly streamlined ecophysiology of SAR11, which relies mostly on passive diffusion from the environment through outer-membrane porins (Giovannoni, 2017), other heterotrophic marine bacteria actively acquire elements using specialised transport systems (Hopkinson & Barbeau, 2012; R. M. Morris et al., 2010; Tang et al., 2012). Our analysis revealed that one of the largest group of proteins significantly enriched in marine BEVs consisted of proteins related to the outer-membrane TonB-dependent transport system (Table S5). The TonB-dependent transport system is a highly diverse functional group of proteins that are involved in cellular uptake of a wide range of substrates, such as iron and iron-containing molecules (e.g., siderophores), as well as vitamins and various organic compounds (Braun, 2024), all of which were observed enriched on BEVs in our dataset.

For BEVs produced by ‘blebbing’ mechanisms, wherein pieces of the outermost cellular membrane detach from the cell to form the vesicle, their proteome is expected to be biased towards outer-membrane proteins (Kulp & Kuehn, 2010). Therefore, in order to determine whether TonB-related proteins are specifically enriched in BEVs, we separately tested the 3,700 outer-membrane proteins identified in our dataset for enrichment. We identified a total of 301 TonB-related proteins that were enriched in BEVs in one or more regions across the transect, compared to 229 TonB-related proteins enriched in the cellular fraction (out of a total 690 and 623 enriched outer-membrane proteins, respectively; Table S6). The majority of the BEV-enriched TonB-dependent proteins belonged to the one-component system of the outer-membrane receptor (OMR) family that perform both binding and transport of the substrate through the outer membrane. The enriched OMRs were primarily affiliated with the orders *Sphingomonadales* (class Alphaproteobacteria), *Alteromonadales* and *Cellvibrionales* (both class Gammaproteobacteria), and more than 20% of them were linked to the utilization of iron (Fig. 8 and Supplementary Text). Along with nitrate, iron was the main limiting micronutrient for surface water primary production across most of the transect (Liu et al., 2024; Yuan et al., 2025). TonB-dependent receptors are highly expressed in heterotrophic bacterial communities in the upper water column (< 100 m depth) (Bergauer et al., 2018). Their strong enrichment on BEVs has previously been observed in cultures of related marine bacteria, including several isolates of *Alteromonadales* (Fadeev et al., 2023), an isolate of endosymbiotic marine *Cellvibrionales* (Gasser et al., 2024), and other marine Gammaproteobacteria (Dürwald et al., 2021; Frias et al., 2010). It has been also observed that iron limitation can lead to enhanced BEVs production in non-marine bacteria (Prados-Rosales et al., 2014) and preferential packaging of iron-related transporters into BEVs has been proposed to facilitate iron accumulation (Lin et al., 2017; Orench-Rivera & Kuehn, 2021; Wang et al., 2021). Taken together, these findings suggest that BEVs might play a role in extracellular scavenging of iron by marine bacteria.

**Fig. 8.**
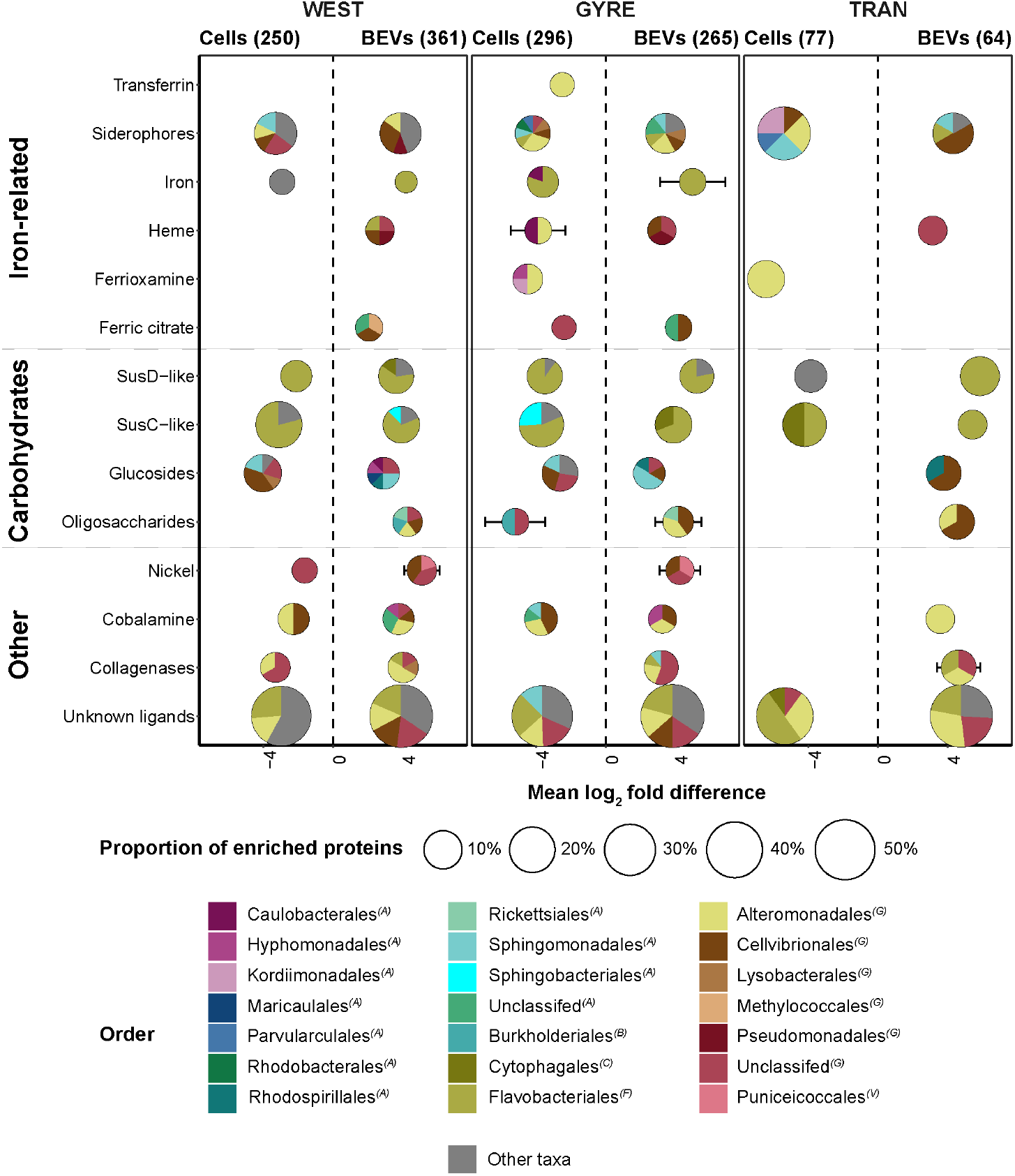
TonB-dependent receptors significantly enriched in BEVs and cellular fractions in different regions. The mean log_2_ fold difference was calculated for each protein group in each fraction. Positive log_2_ fold difference represents significant enrichment in the BEVs fraction, negative log_2_ fold difference represents significant enrichment in the cellular fraction. The total number of enriched proteins in the BEVs and the cellular fractions is provided in brackets. The proteins were grouped according to their predicated ligands. The pie charts show the breakdown of protein taxonomic affiliations on order level within each group. The taxonomic class of each order is given in parentheses: ‘A’ – Alphaproteobacteria, ‘B’ – Betaproteobacteria, ‘C’ – Cytophagia, ‘CY’ – Cyanophyceae, ‘F’ – Flavobacteriia, ‘G’ – Gammaproteobacteria.

To actively transport iron-containing molecules across the outer membrane, such TonB-dependent receptors require proton motive force from the inner membrane (Noinaj et al., 2010). Although BEVs are unlikely to have a proton motive force, these TonB-dependent receptors can still bind the substrates as was recently demonstrated in a culture of a pathogenic Gammaproteobacterium (Dhurve et al., 2022). Taking this into account, we propose that, in conditions of limited availability such as those observed in this study, marine BEVs may ‘collect’ iron in the environment. They could therefore be considered nanoscale ‘hotspots’ of bioavailable iron which could potentially provide a relatively concentrated source for marine bacteria.

### Marine BEVs as mediators of carbohydrate utilisation?

The second largest functional group of TonB-dependent receptors enriched in BEVs was related to the unique SusC-like proteins family (Fig. 8), which play an important role in carbohydrate utilisation by marine Flavobacteria (i.e., order *Flavobacteriales*). Flavobacteria efficiently utilise various sugars using a two component SusCD system, where SusC-like proteins are integral outer-membrane transporters and SusD-like lipoproteins facilitate extracellular high-affinity binding of the sugars (Glenwright et al., 2017). We found a total of 30 SusC-like proteins enriched in BEVs, compared to 55 SusC-like proteins enriched in the cellular fraction. In contrast, there were 27 SusD-like lipoproteins enriched in BEVs, compared to 15 such proteins in the cellular fraction (Table S6). The stronger enrichment of SusD-like lipoproteins in the BEV fraction compared to SusC suggests that flavobacterial BEVs may only be able to bind to carbohydrate molecules, without the ability to transport them. This phenomenon was previously observed in an isolate of marine *Flavobacteriales* (*Formosa* spp.; (Fischer et al., 2019). Trapping carbohydrate molecules on the BEV membrane (or BEV getting attached to a large carbohydrate) may facilitate their efficient digestion by hydrolytic enzymes, in a manner similar to that observed for cellular outer membranes (Reintjes et al., 2017). While we lack such a mechanistic understanding for marine BEVs, the presence of hydrolytic enzymes associated with BEVs and their activity has been observed in several isolates of heterotrophic marine bacteria (Dürwald et al., 2021; Fadeev et al., 2023; J. Li et al., 2016; Naval & Chandra, 2019).

Recent observations have shown that gut bacteria can actively alter the enzyme content of BEVs to optimise carbohydrate utilisation (Sartorio et al., 2023), an analogous process may occur also in marine bacteria. In our dataset, we identified an enrichment of a single glycoside hydrolase (family GH16) in BEVs in the western region (log_2_ fold difference of approximately 4), which was taxonomically affiliated with *Flavobacteriales* (Table S6). The GH16 family contains a diversity of glycoside hydrolases that are active on a wide variety of marine carbohydrate molecules (Viborg et al., 2019). Conceptually, the association of enzymes with marine BEVs is advantageous because it allows them to be delivered to the substrate in a more protected manner compared to being freely dispersed in the water column, and may also allow for more efficient degradation in combination with the SusC-like and SusD like proteins (Du Toit, 2023). When considered alongside other BEV-enriched proteins (Table S6), our observations on marine BEVs are in line with previous findings regarding the composition of BEVs in isolates of marine *Flavobacteriales* and *Alteromonadales* (Dürwald et al., 2021; Fischer et al., 2019). In these studies, it was suggested that BEVs play a role in carbohydrate utilisation, based on their transporter and hydrolytic enzyme content. Previous work has shown that marine bacteria release hydrolytic enzymes into the extracellular milieu to facilitate carbohydrate utilisation during periods of increased phytoplankton activity (Arnosti et al., 2021) - a phenomenon observed in the transitional zone and the western regions of the transect. This suggests that some of the secretion occurs via marine BEVs, potentially affecting degradation rates or access to the products.

### Is the production of marine BEVs a ‘selfish’ trait or a ‘public good’?

The organisms our data indicate to produce most of the BEVs can be found both planktonic (i.e., free-living) and associated with particles (including phytoplankton)(Ivars-Martinez et al., 2008; Seymour et al., 2017). In contrast to the relatively confined particle-associated environments (e.g., biofilms), it is unlikely that BEVs released by a planktonic cell directly into seawater will be encountered again by the same, or a highly similar, bacterial cell (Kashtan et al., 2014; Słomka et al., 2023). This led to the initial hypothesis that marine BEVs likely serve to dispose harmful elements or act as a defence mechanism against viral infection (Biller et al., 2014). However, it has been shown that in some cases the uptake of BEVs only occurs between closely related bacterial lineages (Wang et al., 2021), suggesting that there may be mechanisms that bias interactions between BEVs and recipient cells (Knoke et al., 2020). Thus, BEV production could represent a mechanism through which cells compete for nutrients in the marine environment, potentially favouring only closely related phylogenetic populations.

The environmental observations of natural bacterial communities in this study, alongside previous observations of marine bacterial cultures, may also suggest other scenarios. One alternative explanation is that marine BEVs fulfil their intended function while remaining within the diffusive boundary layer (Stocker & Seymour, 2012). Another possible scenario is that the BEVs remain attached to the cell. Some ‘selfish’ marine bacteria appear to have developed means of producing BEVs that are found in chains attached to the cell. This trait was previously observed in isolates of marine *Alteromonadales* and *Flavobacteriales* (Dürwald et al., 2021; Fischer et al., 2019). Chains of such outer membrane-attached BEVs significantly increase the surface area through which the cell interacts with its environment, facilitating the accumulation of elements necessary for life in the vicinity of the cell and their subsequent uptake. Taking into account the potential stability of marine BEVs for at least a number of days (Biller et al., 2014), they may continue to perform their function (e.g. accumulating phosphate or iron) even after detachment from the cell. Therefore, whether separated from cells or released directly into the environment, marine BEVs could also be considered a ‘public good’ that meets the various needs of the bacterial community.

## Conclusions

Here we have explored BEVs contents within highly diverse natural marine bacterial communities on an extensive oceanic scale in the South Pacific. We found that marine BEVs are ubiquitous in surface seawater and possess a substantial degree of protein diversity. The enrichment of functionally specific proteins in BEVs, which appears to be linked to the ecological conditions experienced by the local bacterial community, which suggests BEVs may contribute to marine biogeochemistry and microbial community metabolism. These findings reinforce the emerging hypothesis that BEVs play a role in nutrient and energy scavenging in the marine environment. Our study provides initial observations on the novel functional potential of marine BEVs, such as their ability to perform local enrichment of molecules necessary for life, which motivate future investigations into the underlying mechanisms. To develop a better understanding of the role of BEVs in the ecophysiology of marine bacteria, research into their uptake dynamics is urgently needed. This is a significant knowledge gap regarding the relationship between BEVs and bacterial cells, which could help us understand whether BEVs are a ‘public good’ or a competitive feature of marine bacterial communities. Based on the extensive oceanographic observations presented in this study, the next stage of the research requires the quantification of the contribution of BEVs to the distribution of essential life-sustaining elements (e.g. iron and phosphate) in the marine environment. This will shed light on their importance for microbial processes in the ocean.

## Materials and methods

### Sample collection

The oceanographic sampling was conducted onboard RV Sonne (cruise SO289 representing section GP21 of the GEOTRACES programme) from 18 February to 8 April 2022 (Fig. 1A). Water samples with a total volume of 60 L were taken from a depth of ca. 6 m at each sampling station using the onboard water pump. To remove particulate matter and eukaryotic cells, water samples were pre-filtered at low pressure using a 3 µm pore size membrane filter (Merck Millipore, MA, USA). From the pre-filtered water sample, 1 mL was preserved with 4% glutaraldehyde (final concentration) and stored at - 80°C until flow cytometric analysis. The pre-filtered seawater sample was then filtered through a 142 mm diameter polycarbonate filter with a pore size of 0.22 µm (Merck Millipore, MA, USA), which were then stored at -80°C and later used for molecular analyses of the bacterial communities. The filtrate (< 0.22 µm) was concentrated ca. 60 times using a Vivaflow tangential flow filtration system (Sartorius, Germany) with a 100 kDa cut-off to 1 L and stored at -20°C until further BEVs purification in the laboratory.

### Phytoplankton pigments analysis

Seawater samples (2-4 L) for diagnostic phytoplankton pigments were filtered onto MF300 glass fibre filters (ThermoFisher Scientific, MA, USA) and stored at -80°C until analysis. Pigments extraction and analysis followed the procedure described by (Browning et al., 2017). Briefly, upon return to a land-based laboratory, pigments were extracted in 90% acetone by homogenization of the filters using glass beads in a cell mill, centrifuged (10 min, 5200 rpm, 4 °C). The supernatant was filtered through 0.2 μm PTFE filters (VWR, PA, USA) and subsequently analysed by reverse-phase high performance liquid chromatography (HPLC) Dionex UltiMate 3000 LC system (ThermoFisher Scientific, MA, USA)(Van Heukelem & Thomas, 2001). Diagnostic pigments were converted to estimated contributions of different phytoplankton types using CHEMTAX (Mackey et al., 1996), with initial pigment ratios according to (DiTullio et al., 2003). Total chlorophyll-a derived from HPLC include both chlorophyll-a and divinyl chlorophyll-a, and is referred to as chlorophyll-a throughout this study.

### Bacterial cell counts

A subsample of 100 μL of seawater from each sampling stations was stained with 1×SYBR Green I (Thermo Fisher Scientific, MA, USA) for 15 min at room temperature and then quantified using FACSAria Flow Cytometer (BD Biosciences, NJ, USA). Counts were processed using R package ‘flowWorkspace’ v4.18.0 (Finak & Jiang, 2022). Quality control of the FCS files was carried out using ‘flow_auto_qc’ function in R package ‘flowAI’ (Monaco et al., 2016). The stained cells were subject to logicle transformation and quantified using the ‘gate_flowclust_2d’ function in R package ‘openCyto’ (Finak et al., 2014), applied on the entire dataset (Fig. S5).

Cell abundances of *Prochlorococcus* and *Synechococcus* were reported by (Yuan et al., 2025) at locations different from those sampled in this study but from the same biogeochemical provinces. Therefore, the abundance of these cells was used to roughly estimate the contribution of cyanobacteria to the total bacterial abundance.

### BEVs isolation and purification

A sub-sample of 100 mL of concentrated seawater was further concentrated about 30 times to a final volume of 3 mL using Vivaspin centrifugal filter units (Sartorius, Germany) with a 100 kDa cut-off. The entire concentrate was then loaded onto an iodixanol density gradient following a published protocol (Biller, Muñoz-Marín, et al., 2022), and centrifuged at 100,000×g at 4°C for 6 h. In the 15-30% and 35-40% fractions, the iodixanol solution was exchanged twice with 1×PBS using Vivaspin centrifugal filter units with a 100 kDa cut-off at 2000×g, and resuspended in 1 mL 1×PBS. From each purified fraction, 800 µL was used for molecular analyses and 200 µL for quantification of BEVs.

### Nanoparticle tracking analysis

The abundance of BEVs was measured by nanoparticle tracking analysis (NTA) using a NanoSight NS300 instrument (Malvern Panalytycal, UK) equipped with a 488 nm laser. Each sample was injected using a syringe pump with a flow speed of 100 μL min^-1^, and five videos of 60 sec were recorded. Videos were analysed using NanoSight NTA software v3.2. All particles in the size up to 300 nm in both the 15-30% and 35-40% fractions were included in the final abundance estimates. The flow cell was thoroughly flushed with Milli-Q water (Merck Millipore) between samples, and visually examined to exclude carry over. The performance of the instrument was routinely checked by measuring the concentration and size distribution of standardized 100 nm silica beads. To estimate the background, we quantified nanoparticles in Optiprep media that was subject to the same washing steps with 1×PBS as the BEVs samples, yielding concentrations of ca. 10^5^ particles mL^-1^, with most particles being below 50 nm in size.

### Metagenomics workflow

One third of the polycarbonate filters with a pore size of 0.22 µm were used for the extraction of genomic DNA using the DNeasy PowerWater Kit (QIAGEN N.V., Germany) according to the manufacturer’s recommendations. The extracted DNA was subject to enzymatic shearing and then shotgun sequencing, performed on two lanes of the NovaSeq S4 platform (Illumina, Inc., CA, USA) at the Vienna Biocenter Core Facilities (https://www.viennabiocenter.org/vbcf/next-generation-sequencing/).

The sequences from all samples were subject to the snakemake (Köster & Rahmann, 2012) metagenomic workflow in Anvi’o v8 (Eren et al., 2021), with recommended parameters. Briefly, the raw FASTQ files were subject to quality control by ‘illumina-utils’ v2.13 (Eren et al., 2013) and taxonomic classification using KrakenUniq v1.0.4 (F. P. Breitwieser et al., 2018) and analysed using R package ‘pavian’ v1.2.1 (Florian P. Breitwieser & Salzberg, 2020). Then the reads were assembled using MEGAHIT v1.2.9 (D. Li et al., 2015) and the gene calling was performed using Prodigal v2.6.3 (Hyatt et al., 2010).

Taxonomic affiliation of the protein coding genes was estimated using Kaiju v1.9.2 (Menzel et al., 2016) against the NCBI RefSeq database (date: 2024-08-14), followed by classification of sequences with unknown taxonomic affiliation against the NCBI nr database (date: 2024-08-25). Functional annotation of the protein coding genes was carried out against InterPro v98 (Blum et al., 2025), NCBIfam v17.0 (Wenjun Li et al., 2021), and Pfam v37 (Mistry et al., 2021) databases using InterProScan v5.66 (Jones et al., 2014). The unidentified proteins sequences with no hits, were assigned to their taxonomy and function using BLASTp (Camacho et al., 2009) best-hit approach against NCBI RefSeq database (date: 2025-01-27). Protein sub-cellular localization predictions were established using DeepLoc v2.0 (Thumuluri et al., 2022). Porins and TonB-dependent receptors were further classified using PSIBLAST (Camacho et al., 2009) against the Transporter Classification Database (TCDB) (Saier et al., 2006), best hit was defined based on identity > 30%, e-value < 0.001 and bit score > 50.

To construct the database for downstream metaproteomic analysis, bacterial protein coding genes were clustered at 99% similarity using CD-HIT v4.6.881 (Weizhong Li & Godzik, 2006).

### Proteomics workflow

We used the remaining two thirds of the 0.22 µm pore-size polycarbonate filters for extraction of cellular proteins. Protein extraction from the filter was performed as described elsewhere (Tinta et al., 2020). Briefly, the filters were ground into small pieces with a sterile metal spatula after submerging the tubes with the filters into liquid nitrogen. Filter pieces were resuspended in lysis buffer (100mM Tris-HCl pH 7.4, 1% SDS, 150mM NaCl, 1mM DTT, 10mM EDTA) and cells were lysed with five freeze-and-thaw cycles. After centrifugation (20,000x *g* at 4°C for 25 min) the supernatant was transferred into a tube and proteins were co-precipitated with 0.015% deoxycholate and 7% trichloroacetic acid (TCA) on ice for 1h and washed twice with ice-cold acetone. Dried protein pellets were resuspended in 50 mM TEAB buffer (Millipore Sigma, Burlington, MA, USA) and cysteines were reduced and alkylated with 10 mM DTT and 55 mM iodoacetamide (IAA), respectively. To extract proteins associated with the BEVs, we added the sample reducing agent NuPAGE (Invitrogen, Waltham, MA, USA) to 600 μL of purified BEVs at 1X final concentration.

All samples were re-precipitated using 9 times the sample volume of 96% EtOH at -20°C overnight. Pellets were resuspended in 50 mM TEAB, followed by overnight in-solution trypsin (Roche, Basel, Switzerland) digestion (1:100, w/w) at 37°C. TFA was added to the samples at 1% final concentration to terminate trypsin digestion. Samples were desalted using Pierce C18 Tips (ThermoFisher Scientific, MA, USA) according to the manufacturer’s protocol. Prior to the LC-MS analysis, digested peptides were dissolved in 0.1% formic acid and 2% acetonitrile.

Purified tryptic peptides were dissolved in 0.1% formic acid (FA) and measured using a nano-reversed-phase high-performance liquid chromatography (RP-HPLC) system Dionex Ultimate 3000 (ThermoFisher Scientific, MA, USA), coupled with a benchtop Quadrupole Orbitrap mass spectrometer Q-Exactive Plus (ThermoFisher Scientific, MA, USA). Peptides were separated on a PepMap RSLC C18 column (ThermoFisher Scientific, MA, USA) at 55°C with a flow rate of 300 nL/min. The two-hour segmented LC gradient ranged from 5% to 80% buffer B (79.9% acetonitrile, 0.1% FA). Mass spectra were acquired in positive ion mode using a top-15 data-dependent acquisition method. Full MS scans were performed at a resolution of 70,000 (*m/z* 200), followed by MS/MS scans at a resolution of 17,500. High-energy collisional dissociation (HCD) fragmentation was applied with a normalized collision energy (NCE) of 30%. Dynamic exclusion was set to 60 s.

The MS raw files of proteins extracted from the bacterial cells and the BEVs were processed using the Spectrum Files RC node in Proteome Discoverer v2.2.0.338 (ThermoFisher Scientific, MA, USA) and searched using SEQUEST-HT against the bacterial protein coding genes from the co-assembled metagenome. Search parameters were as follows: enzyme - trypsin, fragment mass tolerance - 0.8 Da, max. missed cleavages - 2, fixed modifications - carbamidomethyl (Cys), optional modifications - oxidation (Met). Percolator parameters were as follows: max. delta Cn: 0.6, max. rank: 0, validation based on q-value, false discovery rate (calculated by automatic decoy searches) 0.05. Abundance of each protein was estimated by summing the abundance of unique and razor peptides with at least 2 Peptide Spectrum Matches (PSMs). Only proteins with at least one unique peptide and at least two peptides in total (unique and/or razor peptides) were included in the statistical analyses.

### Statistical analyses

Statistical analyses were done in R v4.4.2 (R Core Team 2022) using RStudio v2024.09.1 (RStudio Team 2019). Prior to statistical analyses, the protein abundance matrix was log_2_-transformed and median-centred. Protein enrichment analysis was performed using the R package ‘DEqMS’ v1.22.0 (Y. Zhu, 2018). Only those proteins were included that were observed in at least two samples in each fraction. The criteria for a significantly enriched protein were an absolute log_2_ fold change > 1 and adjusted p-value < 0.1. All figures were generated using the R package ‘ggplot2’ v3.5.1 (Villanueva & Chen, 2019).

## Data availability

The metagenomic raw sequences have been deposited in the European Nucleotide Archive (ENA) at EMBL-EBI under Project accession number PRJEB88044. The mass spectrometry proteomics data have been deposited to the ProteomeXchange Consortium (Deutsch et al., 2023) via the PRIDE (Perez-Riverol et al., 2022) partner repository with the dataset identifier PXD062682.

Scripts for data processing and statistical analyses, as well as raw flow cytometry and NTA files, can be accessed via Zenodo (https://www.zenodo.org/) under doi: 10.5281/zenodo.15772578.

## Author contributions

EF performed fieldwork, the bioinformatic and statistical analyses, and wrote the manuscript. EF and GJH designed the project. EF, NO, TT performed the protein extraction and LAS carried out the LC-MS measurements and wrote the corresponding methods section. HL, ZY and TJB performed the phytoplankton pigment sampling and analyses. DS and SJB contributed to data interpretation and manuscript writing. All authors approved the submitted version of the article.

## Acknowledgments

We thank the captain, the crew and the scientific team of the SO289 research expedition (‘S Pacific GEOTRACES’, funded by the Bundesministerium für Bildung und Forschung (BMBF), Förderkennzeichen 03G0289NA). We thank A. Mutzberg, D. Jasinski, F. Evers, T. Schott, K. Nachtigall and T. Klüver for technical support. We thank Z. Steiner for his support during the expedition. M. Aubert for assistance with laboratory sample processing. EF acknowledges funding by the Austrian Science Fund (FWF) M2797-B. This work was also supported by the Austrian Science Fund (FWF) project I4978-B to GJH and by the Slovenian Research Agency (ARIS) project J7-2599 to TT.

## References

Ahmadian, S., Jafari, N., Tamadon, A., Ghaffarzadeh, A., Rahbarghazi, R., & Mahdipour, M. (2024). Different storage and freezing protocols for extracellular vesicles: a systematic review. Stem Cell Research & Therapy, 15(1), 453.

Angulo-Cánovas, E., Bartual, A., López-Igual, R., Luque, I., Radzinski, N. P., Shilova, I., Anjur-Dietrich, M., García-Jurado, G., Úbeda, B., González-Reyes, J. A., Díez, J., Chisholm, S. W., García-Fernández, J. M., & Del Carmen Muñoz-Marín, M. (2024). Direct interaction between marine cyanobacteria mediated by nanotubes. Science Advances, 10(21), eadj1539.

Arnosti, C., Wietz, M., Brinkhoff, T., Hehemann, J.-H., Probandt, D., Zeugner, L., & Amann, R. (2021). The Biogeochemistry of Marine Polysaccharides: Sources, Inventories, and Bacterial Drivers of the Carbohydrate Cycle. Annual Review of Marine Science, 13, 81–108.

Bar-On, Y. M., & Milo, R. (2019). The Biomass Composition of the Oceans: A Blueprint of Our Blue Planet. Cell, 179(7), 1451–1454.

Bergauer, K., Fernandez-Guerra, A., Garcia, J. A. L., Sprenger, R. R., Stepanauskas, R., Pachiadaki, M. G., Jensen, O. N., & Herndl, G. J. (2018). Organic matter processing by microbial communities throughout the Atlantic water column as revealed by metaproteomics. Proceedings of the National Academy of Sciences of the United States of America, 115(3), E400–E408.

Bianchi, D., Weber, T. S., Kiko, R., & Deutsch, C. (2018). Global niche of marine anaerobic metabolisms expanded by particle microenvironments. Nature Geoscience, 11(4), 263–268.

Biller, S. J., Coe, A., Arellano, A. A., Dooley, K., Silvestri, S. M., Gong, J. S., Yeager, E. A., Becker, J. W., & Chisholm, S. W. (2023). Environmental and Taxonomic Drivers of Bacterial Extracellular Vesicle Production in Marine Ecosystems. Applied and Environmental Microbiology, e0059423.

Biller, S. J., Lundeen, R. A., Hmelo, L. R., Becker, K. W., Arellano, A. A., Dooley, K., Heal, K. R., Carlson, L. T., Van Mooy, B. A. S., Ingalls, A. E., & Chisholm, S. W. (2022). Prochlorococcus extracellular vesicles: molecular composition and adsorption to diverse microbes. Environmental Microbiology, 24(1), 420–435.

Biller, S. J., McDaniel, L. D., Breitbart, M., Rogers, E., Paul, J. H., & Chisholm, S. W. (2017). Membrane vesicles in sea water: heterogeneous DNA content and implications for viral abundance estimates. The ISME Journal, 11(2), 394–404.

Biller, S. J., Muñoz-Marín, M. D. C., Lima, S., Matinha-Cardoso, J., Tamagnini, P., & Oliveira, P. (2022). Isolation and Characterization of Cyanobacterial Extracellular Vesicles. Journal of Visualized Experiments: JoVE, 180. 10.3791/63481

Biller, S. J., Schubotz, F., Roggensack, S. E., Thompson, A. W., Summons, R. E., & Chisholm, S. W. (2014). Bacterial vesicles in marine ecosystems. Science, 343(6167), 183–186.

Bitto, N. J., Chapman, R., Pidot, S., Costin, A., Lo, C., Choi, J., D’Cruze, T., Reynolds, E. C., Dashper, S. G., Turnbull, L., Whitchurch, C. B., Stinear, T. P., Stacey, K. J., & Ferrero, R. L. (2017). Bacterial membrane vesicles transport their DNA cargo into host cells. Scientific Reports, 7(1), 7072.

Blum, M., Andreeva, A., Florentino, L. C., Chuguransky, S. R., Grego, T., Hobbs, E., Pinto, B. L., Orr, A., Paysan-Lafosse, T., Ponamareva, I., Salazar, G. A., Bordin, N., Bork, P., Bridge, A., Colwell, L., Gough, J., Haft, D. H., Letunic, I., Llinares-López, F., … Bateman, A. (2025). InterPro: the protein sequence classification resource in 2025. Nucleic Acids Research, 53(D1), D444–D456.

Bosdriesz, E., Magnúsdóttir, S., Bruggeman, F. J., Teusink, B., & Molenaar, D. (2015). Binding proteins enhance specific uptake rate by increasing the substrate-transporter encounter rate. The FEBS Journal, 282(12), 2394–2407.

Braun, V. (2024). Substrate uptake by TonB-dependent outer membrane transporters. Molecular Microbiology, 122(6), 929–947.

Breitwieser, F. P., Baker, D. N., & Salzberg, S. L. (2018). KrakenUniq: confident and fast metagenomics classification using unique k-mer counts. Genome Biology, 19(1), 198.

Breitwieser, Florian P., & Salzberg, S. L. (2020). Pavian: interactive analysis of metagenomics data for microbiome studies and pathogen identification. Bioinformatics (Oxford, England), 36(4), 1303–1304.

Browning, T. J., Achterberg, E. P., Rapp, I., Engel, A., Bertrand, E. M., Tagliabue, A., & Moore, C. M. (2017). Nutrient co-limitation at the boundary of an oceanic gyre. Nature, 551(7679), 242–246.

Camacho, C., Coulouris, G., Avagyan, V., Ma, N., Papadopoulos, J., Bealer, K., & Madden, T. L. (2009). BLAST+: architecture and applications. BMC Bioinformatics, 10(1), 421.

Capone, D. G., & Hutchins, D. A. (2013). Microbial biogeochemistry of coastal upwelling regimes in a changing ocean. Nature Geoscience, 6(9), 711–717.

Clifton, B. E., Alcolombri, U., Uechi, G.-I., Jackson, C. J., & Laurino, P. (2024). The ultra-high affinity transport proteins of ubiquitous marine bacteria. Nature, 634(8034), 721–728.

Deutsch, E. W., Bandeira, N., Perez-Riverol, Y., Sharma, V., Carver, J. J., Mendoza, L., Kundu, D. J., Wang, S., Bandla, C., Kamatchinathan, S., Hewapathirana, S., Pullman, B. S., Wertz, J., Sun, Z., Kawano, S., Okuda, S., Watanabe, Y., MacLean, B., MacCoss, M. J., … Vizcaíno, J. A. (2023). The ProteomeXchange consortium at 10 years: 2023 update. Nucleic Acids Research, 51(D1), D1539–D1548.

Dhurve, G., Madikonda, A. K., Jagannadham, M. V., & Siddavattam, D. (2022). Outer Membrane Vesicles of Acinetobacter baumannii DS002 Are Selectively Enriched with TonB-Dependent Transporters and Play a Key Role in Iron Acquisition. Microbiology Spectrum, 10(2), e0029322.

DiTullio, G. R., Geesey, M. E., Jones, D. R., Daly, K. L., Campbell, L., & Smith, W.O., Jr. (2003). Phytoplankton assemblage structure and primary productivity along 170°W in the South Pacific Ocean. Marine Ecology Progress Series, 255, 55–80.

Domínguez-Martín, M. A., Sauer, P. V., Kirst, H., Sutter, M., Bína, D., Greber, B. J., Nogales, E., Polívka, T., & Kerfeld, C. A. (2022). Structures of a phycobilisome in light-harvesting and photoprotected states. Nature, 609(7928), 835–845.

Du Toit, A. (2023). Capturing glycans. Nature Reviews. Microbiology, 21(8), 485.

Dürwald, A., Zühlke, M.-K., Schlüter, R., Gebbe, R., Bartosik, D., Unfried, F., Becher, D., & Schweder, T. (2021). Reaching out in anticipation: bacterial membrane extensions represent a permanent investment in polysaccharide sensing and utilization. Environmental Microbiology, 23(6), 3149–3163.

Eren, A. M., Kiefl, E., Shaiber, A., Veseli, I., Miller, S. E., Schechter, M. S., Fink, I., Pan, J. N., Yousef, M., Fogarty, E. C., Trigodet, F., Watson, A. R., Esen, Ö. C., Moore, R. M., Clayssen, Q., Lee, M. D., Kivenson, V., Graham, E. D., Merrill, B. D., … Willis, A. D. (2021). Community-led, integrated, reproducible multi-omics with anvi’o. Nature Microbiology, 6(1), 3–6.

Eren, A. M., Vineis, J. H., Morrison, H. G., & Sogin, M. L. (2013). A filtering method to generate high quality short reads using illumina paired-end technology. PloS One, 8(6), e66643.

Fadeev, E., Carpaneto Bastos, C., Hennenfeind, J. H., Biller, S. J., Sher, D., Wietz, M., & Herndl, G.J. (2023). Characterization of membrane vesicles in Alteromonas macleodii indicates potential roles in their copiotrophic lifestyle. MicroLife, 4, 2022.09.27.509651.

Finak, G., Frelinger, J., Jiang, W., Newell, E. W., Ramey, J., Davis, M. M., Kalams, S. A., De Rosa, S. C., & Gottardo, R. (2014). OpenCyto: an open source infrastructure for scalable, robust, reproducible, and automated, end-to-end flow cytometry data analysis. PLoS Computational Biology, 10(8), e1003806.

Finak, G., & Jiang, M. (2022). flowWorkspace: Infrastructure for representing and interacting with gated and ungated cytometry data sets (4.8.0) [Computer software]. https://www.bioconductor.org/packages/release/bioc/html/flowWorkspace.html

Fischer, T., Schorb, M., Reintjes, G., Kolovou, A., Santarella-Mellwig, R., Markert, S., Rhiel, E., Littmann, S., Becher, D., Schweder, T., & Harder, J. (2019). Biopearling of Interconnected Outer Membrane Vesicle Chains by a Marine Flavobacterium. Applied and Environmental Microbiology, 85(19). 10.1128/AEM.00829-19

Frias, A., Manresa, A., de Oliveira, E., López-Iglesias, C., & Mercade, E. (2010). Membrane vesicles: a common feature in the extracellular matter of cold-adapted antarctic bacteria. Microbial Ecology, 59(3), 476–486.

Gasol, J. M., del Giorgio, P. A., Massana, R., & Duarte, C. M. (1995). Active versus inactive bacteria:size-dependence in a coastal marine plankton community. Marine Ecology Progress Series, 128, 91–97.

Gasser, M. T., Liu, A., Altamia, M. A., Brensinger, B. R., Brewer, S. L., Flatau, R., Hancock, E. R., Preheim, S. P., Filone, C. M., & Distel, D. L. (2024). Membrane vesicles can contribute to cellulose degradation by Teredinibacter turnerae, a cultivable intracellular Endosymbiont of shipworms. Microbial Biotechnology, 17(12), e70064.

Gerritzen, M. J. H., Maas, R. H. W., van den Ijssel, J., van Keulen, L., Martens, D. E., Wijffels, R. H., & Stork, M. (2018). High dissolved oxygen tension triggers outer membrane vesicle formation by Neisseria meningitidis. Microbial Cell Factories, 17(1), 157.

Giovannoni, S. J. (2017). SAR11 Bacteria: The Most Abundant Plankton in the Oceans. Annual Review of Marine Science, 9, 231–255.

Glenwright, A. J., Pothula, K. R., Bhamidimarri, S. P., Chorev, D. S., Baslé, A., Firbank, S. J., Zheng, H., Robinson, C. V., Winterhalter, M., Kleinekathöfer, U., Bolam, D. N., & van den Berg, B. (2017). Structural basis for nutrient acquisition by dominant members of the human gut microbiota. Nature, 541(7637), 407–411.

Hopkinson, B. M., & Barbeau, K. A. (2012). Iron transporters in marine prokaryotic genomes and metagenomes. Environmental Microbiology, 14(1), 114–128.

Hyatt, D., Chen, G.-L., Locascio, P. F., Land, M. L., Larimer, F. W., & Hauser, L. J. (2010). Prodigal: prokaryotic gene recognition and translation initiation site identification. BMC Bioinformatics, 11(1), 119.

Ivars-Martinez, E., Martin-Cuadrado, A.-B., D’auria, G., Mira, A., Ferriera, S., Johnson, J., Friedman, R., & Rodriguez-Valera, F. (2008). Comparative genomics of two ecotypes of the marine planktonic copiotroph Alteromonas macleodii suggests alternative lifestyles associated with different kinds of particulate organic matter. The ISME Journal, 2(12), 1194–1212.

Johnston, E. L., Zavan, L., Bitto, N. J., Petrovski, S., Hill, A. F., & Kaparakis-Liaskos, M. (2023). Planktonic and biofilm-derived Pseudomonas aeruginosa outer membrane vesicles facilitate horizontal gene transfer of Plasmid DNA. Microbiology Spectrum, e0517922.

Jones, P., Binns, D., Chang, H.-Y., Fraser, M., Li, W., McAnulla, C., McWilliam, H., Maslen, J., Mitchell, A., Nuka, G., Pesseat, S., Quinn, A. F., Sangrador-Vegas, A., Scheremetjew, M., Yong, S.-Y., Lopez, R., & Hunter, S. (2014). InterProScan 5: genome-scale protein function classification. Bioinformatics (Oxford, England), 30(9), 1236–1240.

Kadurugamuwa, J. L., & Beveridge, T. J. (1996). Bacteriolytic effect of membrane vesicles from Pseudomonas aeruginosa on other bacteria including pathogens: conceptually new antibiotics. Journal of Bacteriology, 178(10), 2767–2774.

Kamennaya, N. A., Geraki, K., Scanlan, D. J., & Zubkov, M. V. (2020). Accumulation of ambient phosphate into the periplasm of marine bacteria is proton motive force dependent. Nature Communications, 11(1), 2642.

Kashtan, N., Roggensack, S. E., Rodrigue, S., Thompson, J. W., Biller, S. J., Coe, A., Ding, H., Marttinen, P., Malmstrom, R. R., Stocker, R., Follows, M. J., Stepanauskas, R., & Chisholm, S. W. (2014). Single-cell genomics reveals hundreds of coexisting subpopulations in wild Prochlorococcus. Science (New York, N.Y.), 344(6182), 416–420.

Kirchman, D. L. (1994). The uptake of inorganic nutrients by heterotrophic bacteria. Microbial Ecology, 28(2), 255–271.

Knoke, L. R., Abad Herrera, S., Götz, K., Justesen, B. H., Günther Pomorski, T., Fritz, C., Schäkermann, S., Bandow, J. E., & Aktas, M. (2020). Agrobacterium tumefaciens Small Lipoprotein Atu8019 Is Involved in Selective Outer Membrane Vesicle (OMV) Docking to Bacterial Cells. Frontiers in Microbiology, 11, 1228.

Knox, K. W., Vesk, M., & Work, E. (1966). Relation between excreted lipopolysaccharide complexes and surface structures of a lysine-limited culture of Escherichia coli. Journal of Bacteriology, 92(4), 1206–1217.

Koch, A. L. (1996). What size should a bacterium be? A question of scale. Annual Review of Microbiology, 50(1), 317–348.

Köster, J., & Rahmann, S. (2012). Snakemake--a scalable bioinformatics workflow engine. Bioinformatics (Oxford, England), 28(19), 2520–2522.

Kulp, A., & Kuehn, M. J. (2010). Biological functions and biogenesis of secreted bacterial outer membrane vesicles. Annual Review of Microbiology, 64, 163–184.

Li, D., Liu, C.-M., Luo, R., Sadakane, K., & Lam, T.-W. (2015). MEGAHIT: an ultra-fast single-node solution for large and complex metagenomics assembly via succinct de Bruijn graph. Bioinformatics (Oxford, England), 31(10), 1674–1676.

Li, J., Azam, F., & Zhang, S. (2016). Outer membrane vesicles containing signalling molecules and active hydrolytic enzymes released by a coral pathogen Vibrio shilonii AK1. Environmental Microbiology, 18(11), 3850–3866.

Li, Weizhong, & Godzik, A. (2006). Cd-hit: a fast program for clustering and comparing large sets of protein or nucleotide sequences. Bioinformatics (Oxford, England), 22(13), 1658–1659.

Li, Wenjun, O’Neill, K. R., Haft, D. H., DiCuccio, M., Chetvernin, V., Badretdin, A., Coulouris, G., Chitsaz, F., Derbyshire, M. K., Durkin, A. S., Gonzales, N. R., Gwadz, M., Lanczycki, C. J., Song, J. S., Thanki, N., Wang, J., Yamashita, R. A., Yang, M., Zheng, C., … Thibaud-Nissen, F. (2021). RefSeq: expanding the Prokaryotic Genome Annotation Pipeline reach with protein family model curation. Nucleic Acids Research, 49(D1), D1020–D1028.

Li, Z., Clarke, A. J., & Beveridge, T. J. (1998). Gram-negative bacteria produce membrane vesicles which are capable of killing other bacteria. Journal of Bacteriology, 180(20), 5478–5483.

Lin, J., Zhang, W., Cheng, J., Yang, X., Zhu, K., Wang, Y., Wei, G., Qian, P.-Y., Luo, Z.-Q., & Shen, X. (2017). A Pseudomonas T6SS effector recruits PQS-containing outer membrane vesicles for iron acquisition. Nature Communications, 8(1), 14888.

Linney, M. D., Eppley, J. M., Romano, A. E., Luo, E., DeLong, E. F., & Karl, D. M. (2022). Microbial Sources of Exocellular DNA in the Ocean. Applied and Environmental Microbiology, 88(7), e0209321.

Liu, H., Yuan, Z., Gosnell, K. J., Liu, T., Tammen, J. K., Wen, Z., Engel, A., Liu, X., Huang, B., Kao, S.-J., Achterberg, E. P., & Browning, T. J. (2024). Patterns of (micro)nutrient limitation across the South Pacific Ocean. Communications Earth & Environment, 5(1), 1–9.

Lücking, D., Mercier, C., Alarcón-Schumacher, T., & Erdmann, S. (2023). Extracellular vesicles are the main contributor to the non-viral protected extracellular sequence space. ISME Communications, 3(1), 112.

Mackey, M. D., Mackey, D. J., Higgins, H. W., & Wright, S. W. (1996). CHEMTAX - a program for estimating class abundances from chemical markers:application to HPLC measurements of phytoplankton. Marine Ecology Progress Series, 144, 265–283.

Marsili, E., Baron, D. B., Shikhare, I. D., Coursolle, D., Gralnick, J. A., & Bond, D. R. (2008). Shewanella secretes flavins that mediate extracellular electron transfer. Proceedings of the National Academy of Sciences of the United States of America, 105(10), 3968–3973.

Mashburn, L. M., & Whiteley, M. (2005). Membrane vesicles traffic signals and facilitate group activities in a prokaryote. Nature, 437(7057), 422–425.

McMillan, H. M., & Kuehn, M. J. (2021). The extracellular vesicle generation paradox: a bacterial point of view. The EMBO Journal, e108174.

Mella-Flores, D., Six, C., Ratin, M., Partensky, F., Boutte, C., Le Corguillé, G., Marie, D., Blot, N., Gourvil, P., Kolowrat, C., & Garczarek, L. (2012). Prochlorococcus and Synechococcus have evolved different adaptive mechanisms to cope with light and UV stress. Frontiers in Microbiology, 3, 285.

Menzel, P., Ng, K. L., & Krogh, A. (2016). Fast and sensitive taxonomic classification for metagenomics with Kaiju. Nature Communications, 7(1), 11257.

Minhas, T., & Greenman, J. (1989). Production of cell-bound and vesicle-associated trypsin-like protease, alkaline phosphatase and N-acetyl-beta-glucosaminidase by Bacteroides gingivalis strain W50. Journal of General Microbiology, 135(3), 557–564.

Mistry, J., Chuguransky, S., Williams, L., Qureshi, M., Salazar, G. A., Sonnhammer, E. L. L., Tosatto, S. C. E., Paladin, L., Raj, S., Richardson, L. J., Finn, R. D., & Bateman, A. (2021). Pfam: The protein families database in 2021. Nucleic Acids Research, 49(D1), D412–D419.

Monaco, G., Chen, H., Poidinger, M., Chen, J., de Magalhães, J. P., & Larbi, A. (2016). flowAI: automatic and interactive anomaly discerning tools for flow cytometry data. Bioinformatics (Oxford, England), 32(16), 2473–2480.

Moore, C. M., Mills, M. M., Arrigo, K. R., Berman-Frank, I., Bopp, L., Boyd, P. W., Galbraith, E. D., Geider, R. J., Guieu, C., Jaccard, S. L., Jickells, T. D., La Roche, J., Lenton, T. M., Mahowald, N. M., Marañón, E., Marinov, I., Moore, J. K., Nakatsuka, T., Oschlies, A., … Ulloa, O. (2013). Processes and patterns of oceanic nutrient limitation. Nature Geoscience, 6(9), 701–710.

Morel, A., Claustre, H., & Gentili, B. (2010). The most oligotrophic subtropical zones of the global ocean: similarities and differences in terms of chlorophyll and yellow substance. Biogeosciences, 7(10), 3139–3151.

Morris, J. J., Johnson, Z. I., Szul, M. J., Keller, M., & Zinser, E. R. (2011). Dependence of the cyanobacterium Prochlorococcus on hydrogen peroxide scavenging microbes for growth at the ocean’s surface. PloS One, 6(2), e16805.

Morris, R. M., Nunn, B. L., Frazar, C., Goodlett, D. R., Ting, Y. S., & Rocap, G. (2010). Comparative metaproteomics reveals ocean-scale shifts in microbial nutrient utilization and energy transduction. The ISME Journal, 4(5), 673–685.

Morris, R. M., Rappé, M. S., Connon, S. A., Vergin, K. L., Siebold, W. A., Carlson, C. A., & Giovannoni, S. J. (2002). SAR11 clade dominates ocean surface bacterioplankton communities. Nature, 420(6917), 806–810.

Naval, P., & Chandra, T. S. (2019). Characterization of membrane vesicles secreted by seaweed associated bacterium Alteromonas macleodii KS62. Biochemical and Biophysical Research Communications, 514(2), 422–427.

Nekrouf, N. A., Maestre-Carballa, L., Lluesma-Gomez, M., Martinez-Hernandez, F., & Martínez-García, M. (2025). Annual dynamics and metagenomics of marine vesicles: One more layer of complexity in the dissolved organic fraction. Limnology and Oceanography. 10.1002/lno.70031

Noinaj, N., Guillier, M., Barnard, T. J., & Buchanan, S. K. (2010). TonB-dependent transporters: regulation, structure, and function. Annual Review of Microbiology, 64(1), 43–60.

Oggerin, M., Viver, T., Brüwer, J., Voß, D., García-Llorca, M., Zielinski, O., Orellana, L. H., & Fuchs, B. M. (2024). Niche differentiation within bacterial key-taxa in stratified surface waters of the Southern Pacific Gyre. The ISME Journal, 18(1). 10.1093/ismejo/wrae155

Okamoto, A., Saito, K., Inoue, K., Nealson, K. H., Hashimoto, K., & Nakamura, R. (2014). Uptake of self-secreted flavins as bound cofactors for extracellular electron transfer in Geobacter species. Energy & Environmental Science, 7(4), 1357–1361.

Orench-Rivera, N., & Kuehn, M. J. (2016). Environmentally controlled bacterial vesicle-mediated export. Cellular Microbiology, 18(11), 1525–1536.

Orench-Rivera, N., & Kuehn, M. J. (2021). Differential Packaging Into Outer Membrane Vesicles Upon Oxidative Stress Reveals a General Mechanism for Cargo Selectivity. Frontiers in Microbiology, 12, 561863.

Pérez-Cruz, C., Delgado, L., López-Iglesias, C., & Mercade, E. (2015). Outer-inner membrane vesicles naturally secreted by gram-negative pathogenic bacteria. PloS One, 10(1), e0116896.

Perez-Riverol, Y., Bai, J., Bandla, C., García-Seisdedos, D., Hewapathirana, S., Kamatchinathan, S., Kundu, D. J., Prakash, A., Frericks-Zipper, A., Eisenacher, M., Walzer, M., Wang, S., Brazma, A., & Vizcaíno, J. A. (2022). The PRIDE database resources in 2022: a hub for mass spectrometry-based proteomics evidences. Nucleic Acids Research, 50(D1), D543–D552.

Pierella Karlusich, J. J., & Carrillo, N. (2017). Evolution of the acceptor side of photosystem I: ferredoxin, flavodoxin, and ferredoxin-NADP+ oxidoreductase. Photosynthesis Research, 134(3), 235–250.

Pontiller, B., Martínez-García, S., Joglar, V., Amnebrink, D., Pérez-Martínez, C., González, J. M., Lundin, D., Fernández, E., Teira, E., & Pinhassi, J. (2022). Rapid bacterioplankton transcription cascades regulate organic matter utilization during phytoplankton bloom progression in a coastal upwelling system. The ISME Journal, 16(10), 2360–2372.

Prados-Rosales, R., Weinrick, B. C., Piqué, D. G., Jacobs, W. R., Jr, Casadevall, A., & Rodriguez, G. M. (2014). Role for Mycobacterium tuberculosis membrane vesicles in iron acquisition. Journal of Bacteriology, 196(6), 1250–1256.

Reintjes, G., Arnosti, C., Fuchs, B. M., & Amann, R. (2017). An alternative polysaccharide uptake mechanism of marine bacteria. The ISME Journal, 11(7), 1640–1650.

Reintjes, G., Tegetmeyer, H. E., Bürgisser, M., Orlić, S., Tews, I., Zubkov, M., Voß, D., Zielinski, O., Quast, C., Glöckner, F. O., Amann, R., Ferdelman, T. G., & Fuchs, B. M. (2019). On-site analysis of bacterial communities of the ultraoligotrophic South Pacific Gyre. Applied and Environmental Microbiology, 85(14). 10.1128/AEM.00184-19

Renelli, M., Matias, V., Lo, R. Y., & Beveridge, T. J. (2004). DNA-containing membrane vesicles of Pseudomonas aeruginosa PAO1 and their genetic transformation potential. Microbiology (Reading, England), 150(Pt 7), 2161–2169.

Sabra, W., Lünsdorf, H., & Zeng, A.-P. (2003). Alterations in the formation of lipopolysaccharide and membrane vesicles on the surface of Pseudomonas aeruginosa PAO1 under oxygen stress conditions. Microbiology, 149(Pt 10), 2789–2795.

Saier, M. H., Jr, Tran, C.V., & Barabote, R. D. (2006). TCDB: the Transporter Classification Database for membrane transport protein analyses and information. Nucleic Acids Research, 34(Database issue), D181–6.

Sartorio, M. G., Pardue, E. J., Scott, N. E., & Feldman, M. F. (2023). Human gut bacteria tailor extracellular vesicle cargo for the breakdown of diet- and host-derived glycans. Proceedings of the National Academy of Sciences of the United States of America, 120(27), e2306314120.

Scanlan, D. J., Ostrowski, M., Mazard, S., Dufresne, A., Garczarek, L., Hess, W. R., Post, A. F., Hagemann, M., Paulsen, I., & Partensky, F. (2009). Ecological genomics of marine picocyanobacteria. Microbiology and Molecular Biology Reviews: MMBR, 73(2), 249–299.

Schwechheimer, C., & Kuehn, M. J. (2015). Outer-membrane vesicles from Gram-negative bacteria: biogenesis and functions. Nature Reviews. Microbiology, 13(10), 605–619.

Schwechheimer, C., Sullivan, C. J., & Kuehn, M. J. (2013). Envelope control of outer membrane vesicle production in Gram-negative bacteria. Biochemistry, 52(18), 3031–3040.

Seymour, J. R., Amin, S. A., Raina, J.-B., & Stocker, R. (2017). Zooming in on the phycosphere: the ecological interface for phytoplankton–bacteria relationships. Nature Microbiology, 2(7), 1– 12.

Słomka, J., Alcolombri, U., Carrara, F., Foffi, R., Peaudecerf, F. J., Zbinden, M., & Stocker, R. (2023). Encounter rates prime interactions between microorganisms. Interface Focus, 13(2), 20220059.

Soler, N., & Forterre, P. (2020). Vesiduction: the fourth way of HGT. Environmental Microbiology, 22(7), 2457–2460.

Soler, N., Krupovic, M., Marguet, E., & Forterre, P. (2015). Membrane vesicles in natural environments: a major challenge in viral ecology [Review of Membrane vesicles in natural environments: a major challenge in viral ecology]. The ISME Journal, 9(4), 793–796. nature.com.

Sowell, S. M., Wilhelm, L. J., Norbeck, A. D., Lipton, M. S., Nicora, C. D., Barofsky, D. F., Carlson, C. A., Smith, R. D., & Giovanonni, S. J. (2009). Transport functions dominate the SAR11 metaproteome at low-nutrient extremes in the Sargasso Sea. The ISME Journal, 3(1), 93–105.

Stocker, R. (2012). Marine microbes see a sea of gradients. Science, 338(6107), 628–633.

Stocker, R., & Seymour, J. R. (2012). Ecology and physics of bacterial chemotaxis in the ocean. Microbiology and Molecular Biology Reviews, 76(4), 792–812.

Tang, K., Jiao, N., Liu, K., Zhang, Y., & Li, S. (2012). Distribution and functions of TonB-dependent transporters in marine bacteria and environments: implications for dissolved organic matter utilization. PloS One, 7(7), e41204.

Teeling, H., Fuchs, B. M., Bennke, C. M., Krüger, K., Chafee, M., Kappelmann, L., Reintjes, G., Waldmann, J., Quast, C., Glöckner, F. O., Lucas, J., Wichels, A., Gerdts, G., Wiltshire, K. H., & Amann, R. I. (2016). Recurring patterns in bacterioplankton dynamics during coastal spring algae blooms. ELife, 5, e11888.

Thumuluri, V., Almagro Armenteros, J. J., Johansen, A. R., Nielsen, H., & Winther, O. (2022). DeepLoc 2.0: multi-label subcellular localization prediction using protein language models. Nucleic Acids Research, 50(W1), W228–W234.

Tinta, T., Zhao, Z., Escobar, A., Klun, K., Bayer, B., Amano, C., Bamonti, L., & Herndl, G. J. (2020). Microbial Processing of Jellyfish Detritus in the Ocean. Frontiers in Microbiology, 11, 590995.

Tortell, P. D., Maldonado, M. T., Granger, J., & Price, N. M. (1999). Marine bacteria and biogeochemical cycling of iron in the oceans. FEMS Microbiology Ecology, 29(1), 1–11.

Toyofuku, M., Morinaga, K., Hashimoto, Y., Uhl, J., Shimamura, H., Inaba, H., Schmitt-Kopplin, P., Eberl, L., & Nomura, N. (2017). Membrane vesicle-mediated bacterial communication. The ISME Journal, 11(6), 1504–1509.

Toyofuku, M., Schild, S., Kaparakis-Liaskos, M., & Eberl, L. (2023). Composition and functions of bacterial membrane vesicles. Nature Reviews. Microbiology. 10.1038/s41579-023-00875-5

Toyofuku, M., Tashiro, Y., Hasegawa, Y., Kurosawa, M., & Nomura, N. (2015). Bacterial membrane vesicles, an overlooked environmental colloid: Biology, environmental perspectives and applications. Advances in Colloid and Interface Science, 226(Pt A), 65–77.

Van Heukelem, L., & Thomas, C. S. (2001). Computer-assisted high-performance liquid chromatography method development with applications to the isolation and analysis of phytoplankton pigments. Journal of Chromatography A, 910(1), 31–49.

Viborg, A. H., Terrapon, N., Lombard, V., Michel, G., Czjzek, M., Henrissat, B., & Brumer, H. (2019). A subfamily roadmap of the evolutionarily diverse glycoside hydrolase family 16 (GH16). The Journal of Biological Chemistry, 294(44), 15973–15986.

Villanueva, R. A. M., & Chen, Z. J. (2019). ggplot2: Elegant Graphics for Data Analysis (2nd ed.).c Measurement: Interdisciplinary Research and Perspectives, 17(3), 160–167.

Wang, M., Nie, Y., & Wu, X.-L. (2021). Membrane vesicles from a Dietzia bacterium containing multiple cargoes and their roles in iron delivery. Environmental Microbiology, 23(2), 1009– 1019.

Worden, A. Z., Follows, M. J., Giovannoni, S. J., Wilken, S., Zimmerman, A. E., & Keeling, P. J. (2015). Environmental science. Rethinking the marine carbon cycle: factoring in the multifarious lifestyles of microbes. Science (New York, N.Y.), 347(6223), 1257594.

Yaron, S., Kolling, G. L., Simon, L., & Matthews, K. R. (2000). Vesicle-mediated transfer of virulence genes from Escherichia coli O157:H7 to other enteric bacteria. Applied and Environmental Microbiology, 66(10), 4414–4420.

Yonezawa, H., Osaki, T., Kurata, S., Fukuda, M., Kawakami, H., Ochiai, K., Hanawa, T., & Kamiya, S. (2009). Outer membrane vesicles of Helicobacter pylori TK1402 are involved in biofilm formation. BMC Microbiology, 9, 197.

Yuan, Z., Liu, H., Gosnell, K. J., Wen, Z., Engel, A., Dai, M., Achterberg, E. P., & Browning, T. J. (2025). Localized nutrient colimitation of phytoplankton growth rates across the subtropical South Pacific Ocean. Proceedings of the National Academy of Sciences of the United States of America, 122(50), e2526930122.

Zhang, Z., Xu, A., Hathorne, E., Gutjahr, M., Browning, T., Gosnell, K. J., Liu, T., Steiner, Z., Kiko, R., Yuan, Z.-X., Liu, H., Achterberg, E. P., & Frank, M. (2024). Substantial trace metal input from the 2022 Hunga Tonga-Hunga Ha’apai eruption into the South Pacific. Nature Communications, 15(1), 8986.

Zhu, W.-J., Wang, C., Liu, L., Li, J.-X., Wang, H.-Q., Wang, M.-Q., Cao, H.-Y., Chen, X.-L., Qin, Q.-L., Zhang, Y.-Z., Sun, M.-L., & Wang, P. (2025). Structural and molecular basis for phosphate recognition by SAR11 bacteria. MBio, 16(9), e0165425.

Zhu, Y. (2018). DEqMS. Bioconductor. 10.18129/B9.BIOC.DEQMS

Zlatkov, N., Nadeem, A., Uhlin, B. E., & Wai, S. N. (2020). Eco-evolutionary feedbacks mediated by bacterial membrane vesicles. FEMS Microbiology Reviews. 10.1093/femsre/fuaa047

Zoccarato, L., Sher, D., Miki, T., Segrè, D., & Grossart, H.-P. (2022). A comparative whole-genome approach identifies bacterial traits for marine microbial interactions. Communications Biology, 5(1), 276.

