## Supplementary Text and Figures for "Bacterial extracellular vesicles exhibit distinct functional potential across the biogeographic provinces of the South Pacific Ocean"

To infer the potential ligands of the TonB-dependent receptors we performed homology comparison to previously characterized and manually curated OMR proteins (see Materials and Methods) and found that both in BEVs and cellular fractions ca. 20% of the enriched TonB-dependent receptors were linked to iron utilization (a total of 76 and 52 proteins, respectively). Interestingly, we identified only two TonB-dependent receptors for $Fe^{3+}$, a rapidly depleted dissolved inorganic species of iron, that were enriched in BEVs, while six such receptors were found enriched in the cellular fraction, mostly affiliated with *Flavobacteriales* (Fig. 8). On contrast, we found six different TonB-dependent receptors for heme, an important intracellular iron-binding molecule, that were significantly enriched in BEVs across the different regions, and were taxonomically affiliated with different lineages within the Gammaproteobacteria. Two of such receptors were also enriched in the cellular fraction (Fig. 8). We further identified nine different TonB-dependent receptors of ferric citrate, a metabolic byproduct that can bind iron, significantly enriched in BEVs, while only two such proteins were found enriched in the cellular fraction (Fig. 8). These iron-binding molecules can enter the environment through cell destruction processes, such as viral lysis and grazing (Gledhill, 2007). They can account for a significant proportion of the biogenic iron present in surface waters (Louropoulou et al., 2020) and can be utilized by various marine heterotrophic bacteria using TonB-dependent receptors (Hogle et al., 2017; Hopkinson et al., 2008; Toulza et al., 2012).

Some heterotrophic marine bacteria also perform active scavenging for iron by releasing siderophores (*i.e.*, ferric chelates). These are small molecules that bind strongly iron in the environment and are subsequently acquired by bacterial cells through specialised TonB-dependent receptors (Armstrong et al., 2004; Hogle et al., 2022; Hopkinson & Barbeau, 2012). Production of siderophores in marine bacterial communities is mostly restricted to the class Gammaproteobacteria, however, the capacity to acquire them from the environment (i.e., synthesize specific TonB-dependent receptors) is a taxonomically widespread functional trait (Zoccarato et al., 2022). In our dataset we identified a total of 52 TonB-dependent receptors for various siderophore molecules enriched in BEVs, compared to 35 such receptors enriched in the cellular fraction (Fig. 8). The vast majority of these receptors were taxonomically assigned to the orders *Alteromonadales* and *Cellvibrionales* (both in the class Gammaproteobacteria). While little is known about the ecology and the physiology of *Cellvibrionales*, they are often observed during elevated primary production conditions (e.g., (Pontiller et al., 2022, 2021). Marine *Alteromonadales*, on the other hand, are well-studied copiotrophs that also thrive under high availability of organic matter (Wietz et al., 2022) and are considered to be major siderophore producers and key mediators of iron acquisition in marine bacterial communities (Manck et al., 2020, 2022). Taking this into account, we propose that during episodes of high metabolic activity (e.g. associated with increased primary production), some marine bacteria, such as *Alteromonadales,* not only release siderophores that bind dissolved iron, but also produce BEVs that may 'concentrate' them in the environment.

### Supplementary Figures

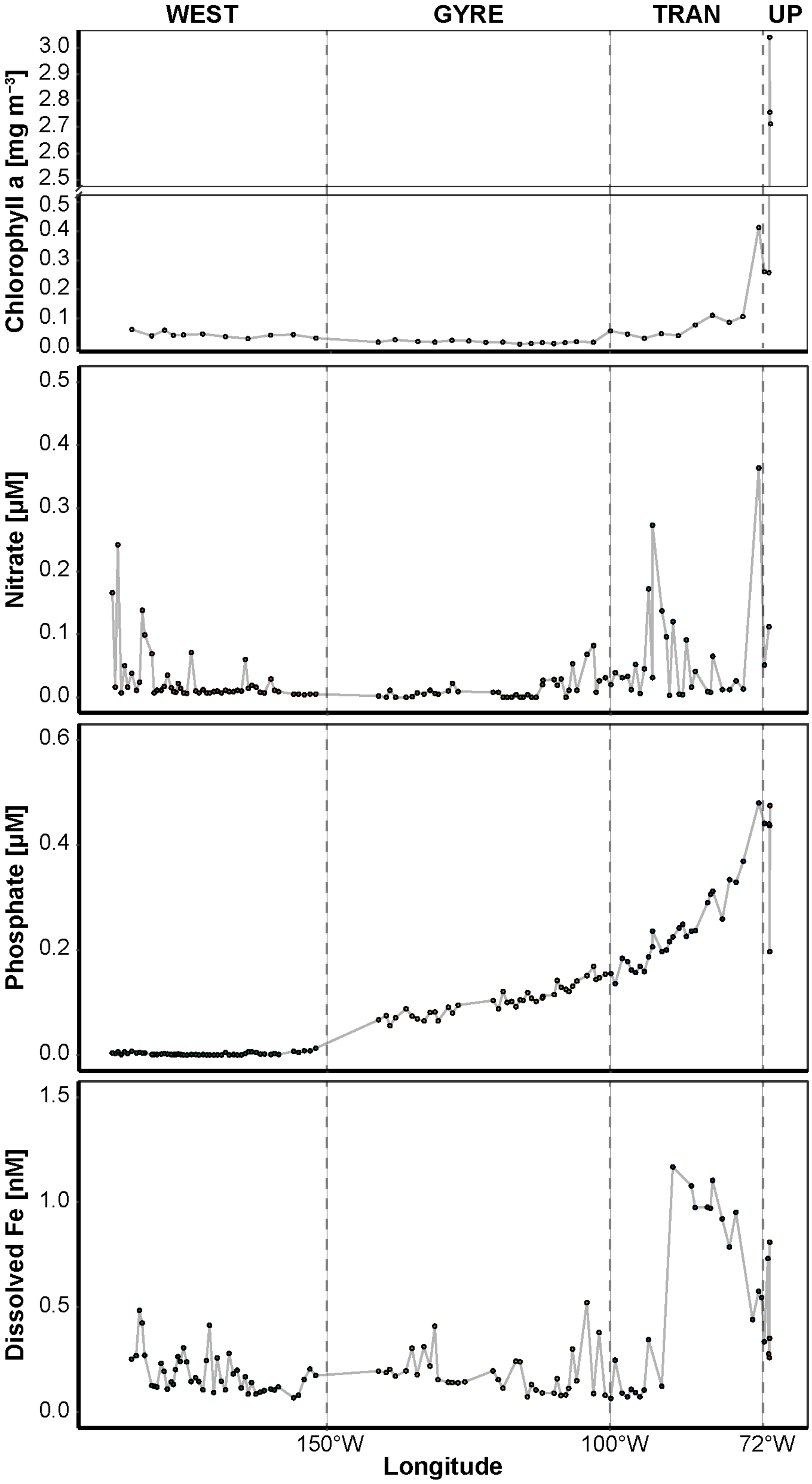


***Fig. S1 – Surface water biogeochemical characteristics across the sampling transect.*** *Spatial grouping of the sampling stations according to four oceanic provinces: 'UP' - Chilean coastal upwelling zone; 'TRAN' - transition zone between the upwelling and the subtropical gyre; 'GYRE' - South Pacific subtropical gyre; 'WEST' - westernmost region. Please note the scale differences on y axis. Data were extracted from Liu et al. (2024).*

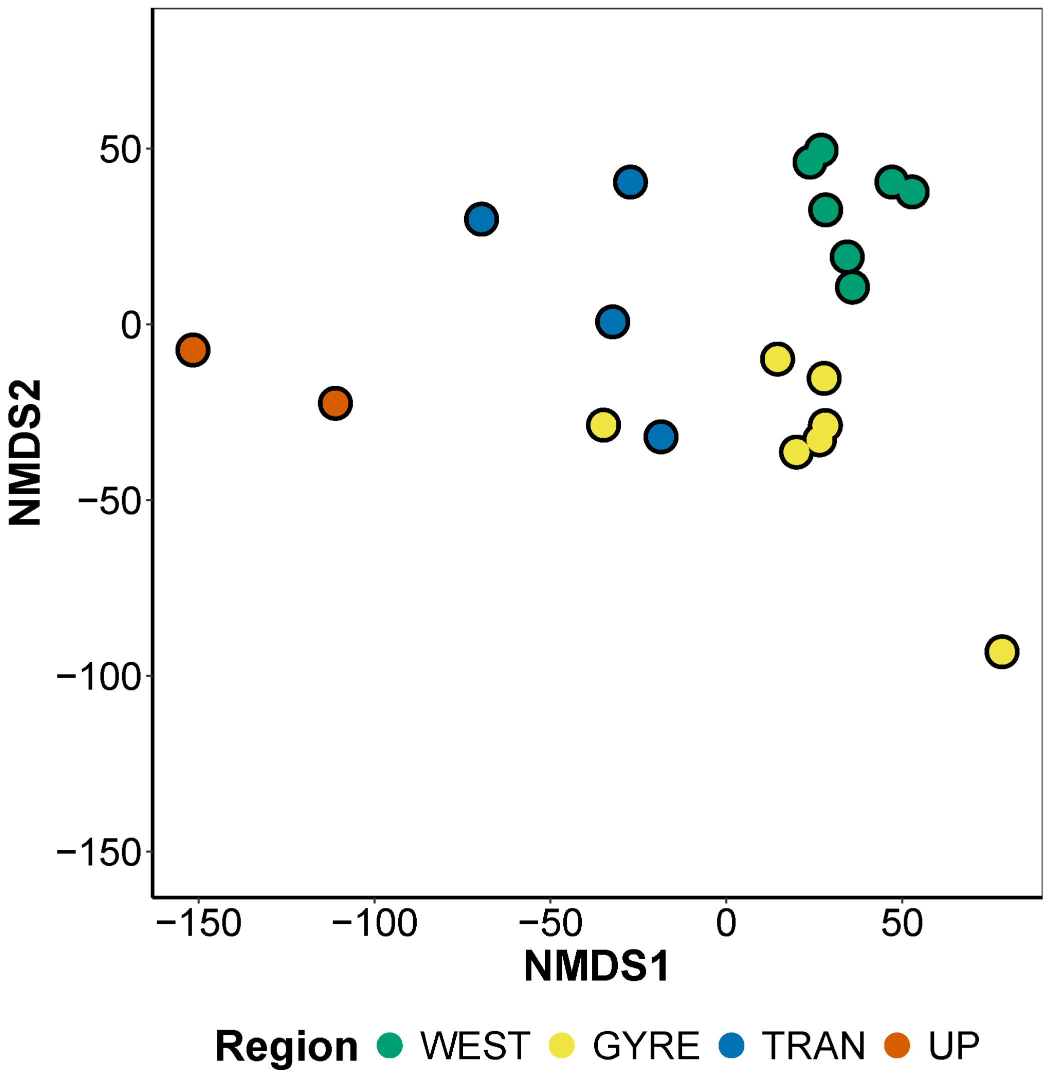


***Fig. S2 – Non-metric Multidimensional Scaling (NMDS) dissimilarities plot of cellular proteomes in each station****. Colours represent the different regions. Dissimilarity was calculated using the Euclidean distance matrix based on log-transformed protein abundances.*

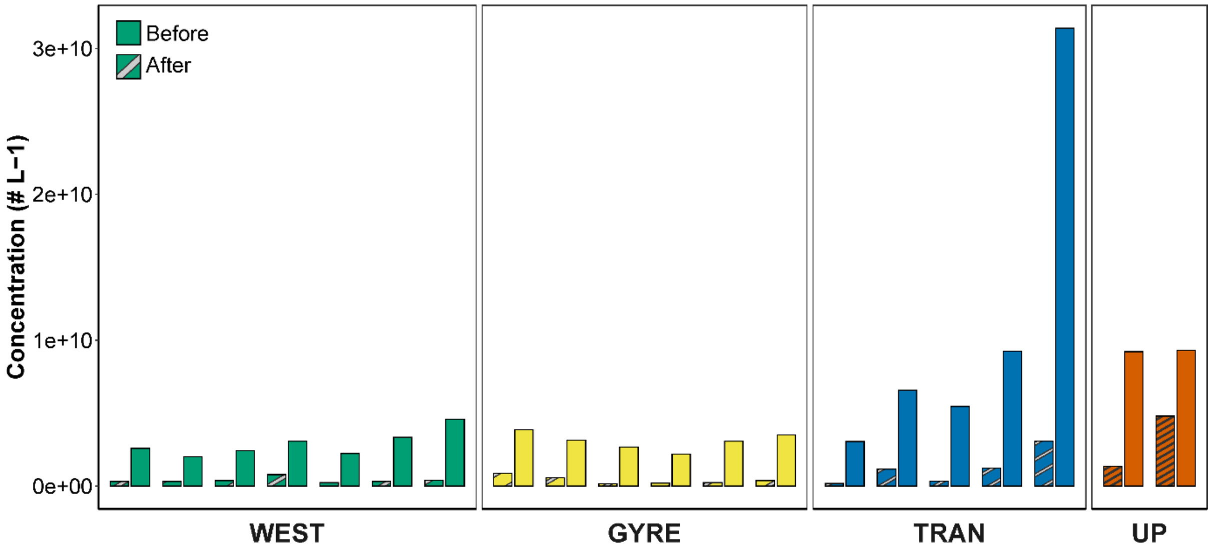


***Fig. S3 – Estimated concentrations of nanoparticles in the size range of 100 kDa-0.22 µm before and after density gradient purification.*** *The nanoparticles before purification are likely comprised of extracellular vesicles, viral-particles and other colloidal material. Whereas after purification the nanoparticles comprise mostly of BEV-like structures.*

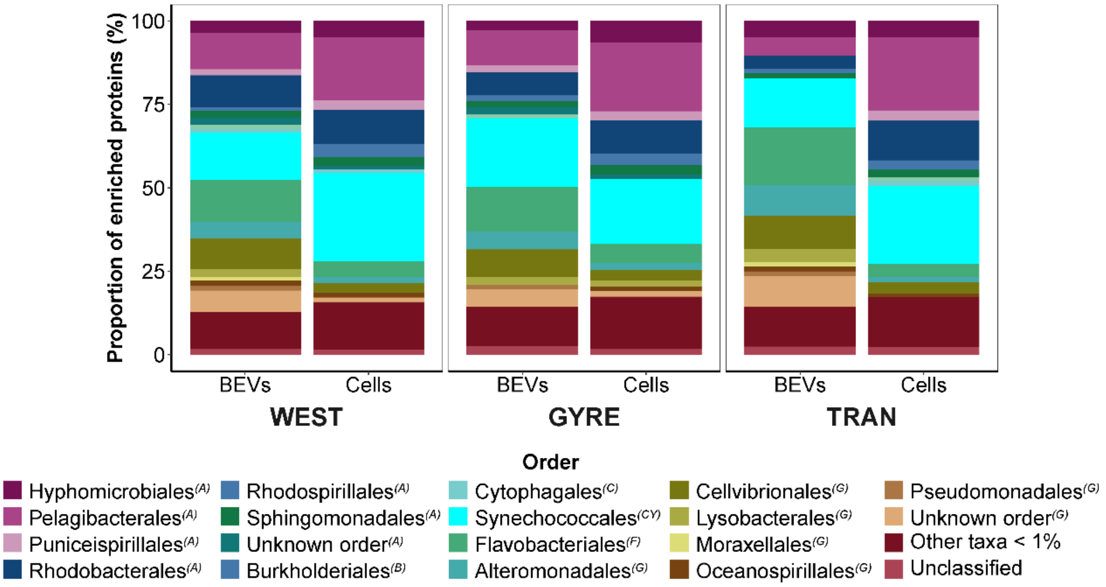


***Fig. S4*** *– Taxonomic affiliations of enriched proteins in the cellular and BEVs fractions according to different oceanic provinces. The taxonomic class of each order is given in parentheses: 'A' – Alphaproteobacteria, 'B' – Betaproteobacteria, 'C' – Cytophagia, 'CY' – Cyanophyceae, 'F' – Flavobacteriia, 'G' – Gammaproteobacteria.*

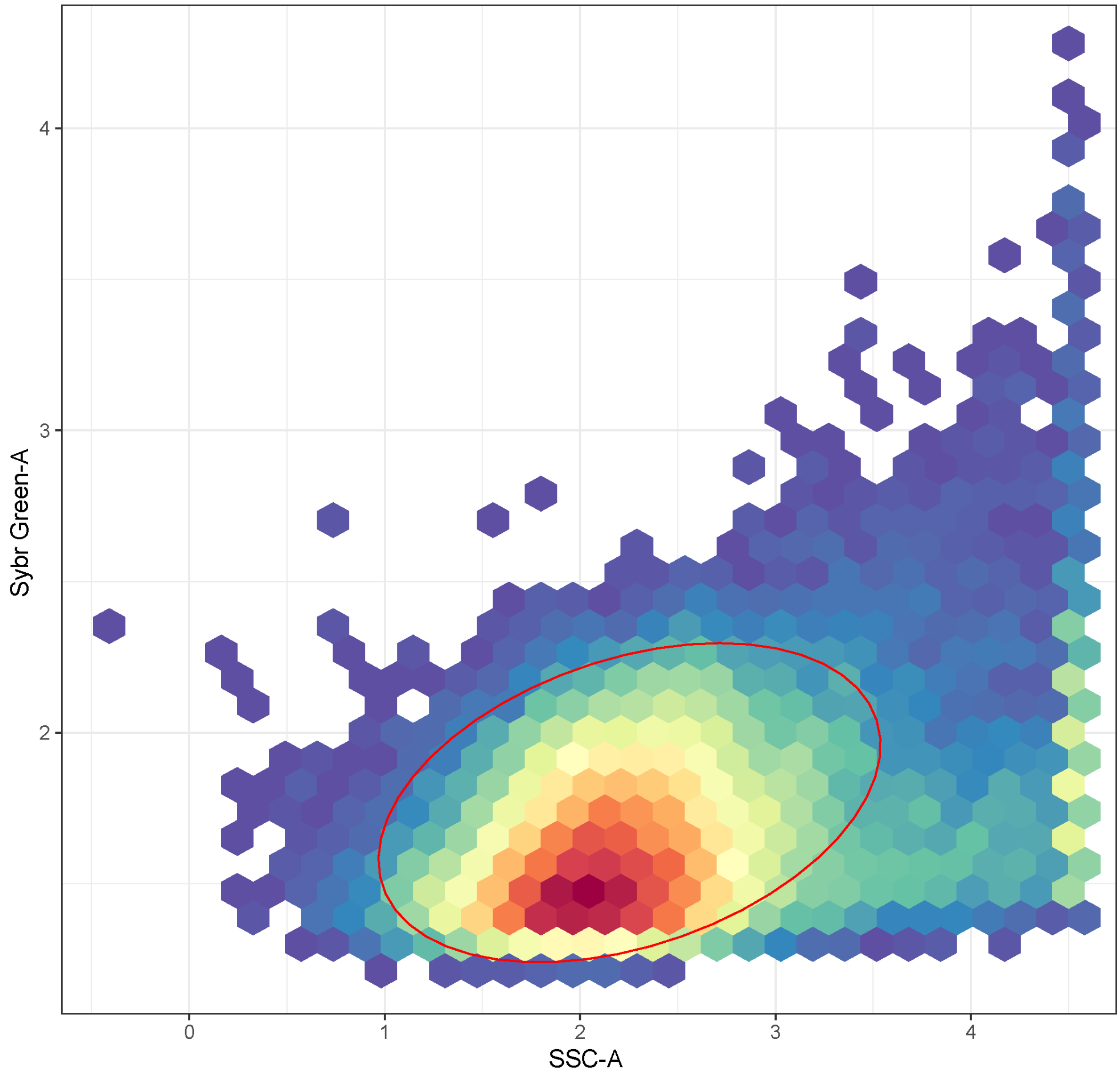


**Fig. S5** – Representation of the gating strategy for cell counts using 'gate_flowclust_2d' function in R package 'openCyto' applied on the entire dataset. The red ellipse shows the area in which the abundance of bacteria was quantified.
